# Lateral prefrontal cortex controls interplay between working memory and actions

**DOI:** 10.1101/2024.09.17.613601

**Authors:** Anastasia Kiyonaga, Jacob A. Miller, Mark D’Esposito

## Abstract

Humans must often keep multiple task goals in mind, at different levels of priority and immediacy, while also interacting with the environment. We might need to remember information for an upcoming task while engaged in more immediate actions. Consequently, actively maintained working memory (WM) content may bleed into ongoing but unrelated motor behavior. Here, we experimentally test the impact of WM maintenance on action execution, and we transcranially stimulate lateral prefrontal cortex (PFC) to parse its functional contributions to WM-motor interactions. We first created a task scenario wherein human participants (both sexes) executed cued hand movements during WM maintenance. We manipulated the compatibility between WM and movement goals at the trial level and the statistical likelihood that the two would be compatible at the block level. We found that remembering directional words (e.g., ‘left’, ‘down’) biased the trajectory and speed of hand movements that occurred during the WM delay, but the bias was dampened in blocks when WM content predictably conflicted with movement goals. Then we targeted left lateral PFC with two different transcranial magnetic stimulation (TMS) protocols before participants completed the task. We found that an intermittent theta-burst protocol, which is thought to be excitatory, dampened sensitivity to block-level control demands (i.e., proactive control), while a continuous theta-burst protocol, which is thought to be inhibitory, dampened adaptation to trial-by-trial conflict (i.e., reactive control). Therefore, lateral PFC is involved in controlling the interplay between WM content and manual action, but different PFC mechanisms may support different time-scales of adaptive control.

**Significance Statement:** Working memory (WM) allows us to keep information active in mind to achieve our moment-to-moment goals. However, WM maintenance may sometimes unintentionally shape our externally-geared actions. This study formalizes the everyday “action slips” humans commit when we type out or say the wrong word in conversation because it was held in mind for a different goal. The results show that internally maintained content can influence ongoing hand movements, but this interplay between WM and motor behavior depends on the cortical excitability state of the lateral PFC. Neural perturbation with transcranial magnetic stimulation (TMS) shows that temporarily increasing or decreasing PFC excitability can make participants more or less susceptible to the impact of WM on actions.

---

Actions sometimes come out differently than planned. If motor intentions are at odds with ongoing thoughts, behavior can be skewed. For instance, we might accidentally type out or speak the wrong word aloud if it is going through our mind. Similar ‘action slips’ are exacerbated in patients with frontal cortex damage, who tend to execute behaviors that are inappropriate for the context (Schwartz, 1995). In typical function, such slips may stem from the adaptive role of working memory (WM) in guiding goal-directed behavior. A tight linkage between WM maintenance and motor execution would keep action plans aligned with top-down goals, but might sometimes allow WM content to “leak” into ongoing behavior. Here, we experimentally formalize everyday action slips, and we examine the neural circuitry that modulates the interplay between WM and motor behavior.

Although WM is often construed as retrospective storage for past information, its core function may be to guide upcoming behavior (Fuster, 2001; Jin et al., 2020; Olivers & Roelfsema, 2020; Postle, 2006; Theeuwes et al., 2009; van Ede et al., 2019). WM activation might signal that the maintained information is pertinent to current goals, explaining why WM content often captures or biases visual attention (Soto et al., 2005, 2008; Soto & Humphreys, 2007). In a prior behavioral study, we found that WM content can also bias hand movements (Miller et al., 2020). During the WM maintenance period for directional word stimuli (e.g., remember ‘left’, ‘down’), cued movements were skewed in a WM-matching direction. Movements were therefore improved if they coincided with WM content but impaired if they conflicted, suggesting that WM may be akin to action preparation (Olivers & Roelfsema, 2020). However, this movement effect was also dampened in task blocks when WM information was less likely to align with movement goals. The interplay between WM content and ongoing motor behavior is therefore adjustable to changing goal states, suggesting that occasional action slips emerge from a generally adaptive system (Carlisle & Woodman, 2011; Kiyonaga et al., 2012; Olivers et al., 2011). This convergence and control between WM and motor behavior could stem from many stages of processing, however, and its source remains to be identified.

Lateral prefrontal cortex (PFC) plays a key role in cortico-striatal circuitry for selecting and using WM content to guide behavior (Chatham et al., 2014). The region is thought to monitor and resolve stimulus-response conflict in tasks like the Stroop or Flanker (Botvinick et al., 2001), and WM content may induce similar conflict (Kiyonaga & Egner, 2014; Pan et al., 2019; Wang et al., 2021). The mid-lateral PFC is also more broadly considered a nexus of cognitive coordination and control (Badre & Nee, 2018; Nee & D’Esposito, 2016). Lateral PFC is a likely candidate to control the interplay between WM and actions, but PFC is multidimensional — it is engaged by a variety of different task demands, shows diverse connectivity, and PFC neurons can exhibit selectivity for various features (Duncan, 2010; Rigotti et al., 2013; Wang et al., 2023). Likewise, there are competing ideas about the role that human PFC plays in WM and attentional control processes (Curtis & D’Esposito, 2003; Leavitt et al., 2017), and PFC regions may implement distinct forms of control through different circuit mechanisms (Braver, 2012).

Here, we ask how lateral PFC contributes to the interplay between WM and concurrent motor behavior. In a dual-task scenario, subjects executed cued hand movements during WM maintenance, and we manipulated the compatibility between WM and action goals from trial-to-trial, as well as the proportion of compatible trials across task blocks. We replicated prior behavioral findings in a baseline condition, and we stimulated left lateral PFC with two transcranial magnetic stimulation (TMS) protocols in experimental comparison conditions. We compared the effects of continuous vs. intermittent theta-burst stimulation (cTBS vs. iTBS), which are thought to have distinct effects on cortical excitability (Huang et al., 2005), subcortical dopamine release (Aceves-Serrano et al., 2022), and network propagation (Cocchi et al., 2015). Whereas cTBS is associated with cortical inhibition, iTBS is associated with excitation. These distinct protocols, delivered to the same region, should therefore serve as strong active controls for one another, to differentiate between functions that rely on the stimulated region. For instance, selective TMS effects on WM performance would implicate the target region in WM storage, while modulation of WM-action compatibility effects would illuminate its role in adaptive control. Moreover, different TMS effects on block-level vs. trial-level indices would illustrate the PFC contribution to superordinate (proactive) task control vs. adapting to immediate changes in (reactive) cognitive control demand. This work will inform the underlying functions that promote WM-driven action slips, clarify the role of PFC in this everyday behavior, and illuminate the cognitive and neural conditions under which these flukes are most likely to occur.

## Materials and Methods

### Overview of experimental procedures

On Day 1, participants underwent an anatomical MRI followed by 10 minutes of a resting-state functional MRI. The anatomical MRI was used to stereotactically navigate the TMS coil over the left lateral PFC target. We targeted a PFC region considered part of a cortico-striatal pathway for the selection and use of WM (Chatham et al., 2014; Strafella et al., 2001). After scanning, participants underwent TMS motor thresholding to calibrate the stimulation intensity for TBS sessions. On Days 2-4, participants received either cTBS, iTBS, or a no-TMS baseline condition, and then they completed the behavioral task (**Fig. 1a**). The order of stimulation conditions was counter-balanced across participants.

**Figure 1.**
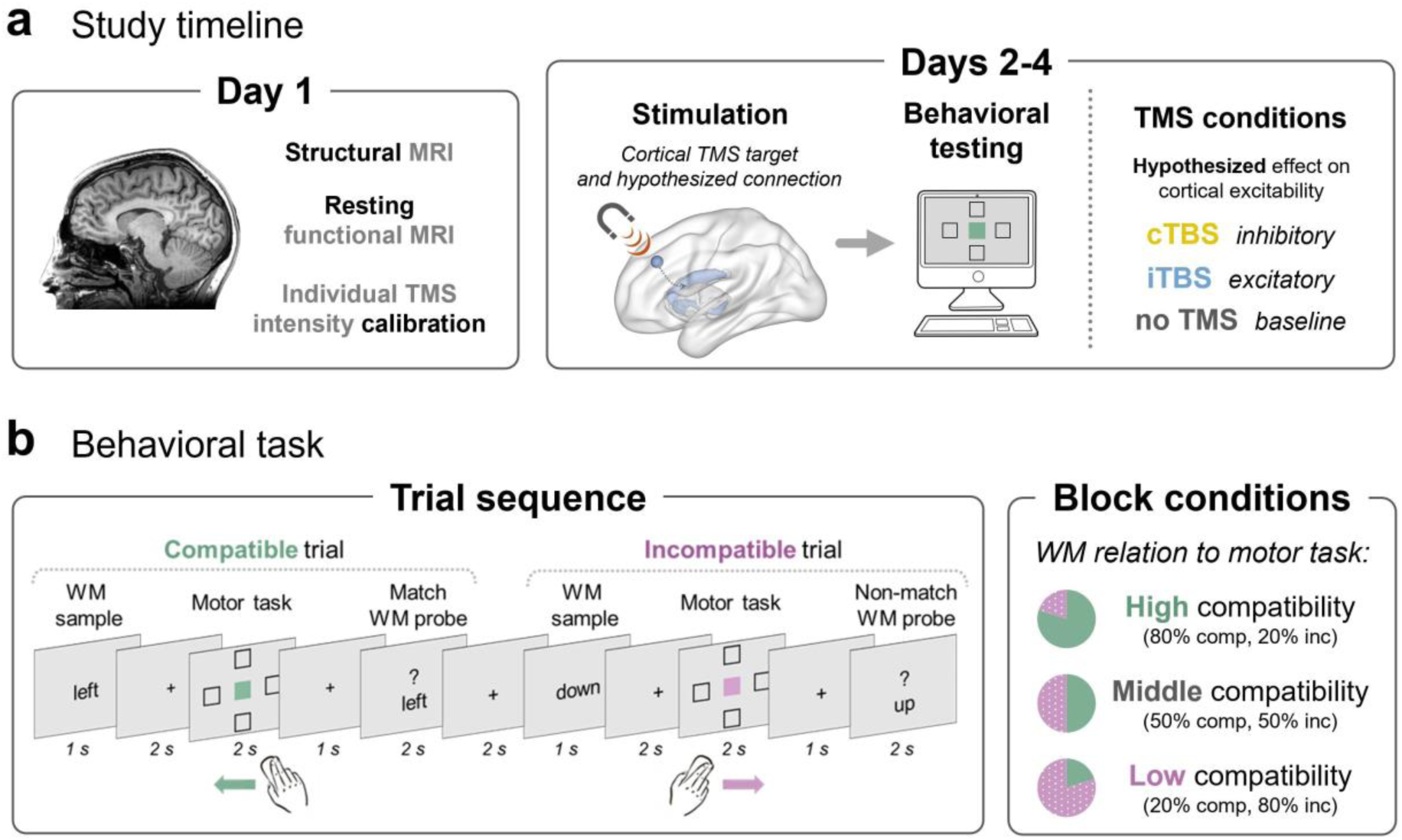
Overview of experimental procedures. (a) On Day 1, participants underwent anatomical and resting-state functional MRI for TMS neuronavigation, then TMS motor thresholding to calibrate their experimental stimulation intensity. On Days 2-4, participants received either a no-TMS baseline condition or one of two experimental TMS protocols to the lateral PFC target, followed by the behavioral task. TMS conditions were given in counterbalanced order on different days. (b) On each trial, participants were first shown a WM sample word to remember (‘up,’ ‘down,’ ‘left,’ or ‘right’), then were cued to click one of four screen locations. They were then given a same/different recognition memory test for the sample word. The meaning of the verbal WM content could be either compatible (green) or incompatible (purple) with the direction of the intervening mouse movement action. The proportion of compatible to incompatible trials was manipulated across blocks of the experiment.

As in our previous behavioral study (Miller et al., 2020), we used a verbal delayed recognition task, interleaved with a simple motor task during the delay period (**Fig. 1b**). On each trial, participants were instructed to remember a directional WM sample word (‘up,’ ‘down,’ ‘left,’ or ‘right’). They were then visually cued to click a screen location (to the top, bottom, left, or right of center). After the cued mouse click, participants were tested on their memory for the sample word. On each trial, we manipulated whether the meaning of the verbal WM content was either compatible (e.g., remember ‘left’, click inside leftward box) or incompatible (e.g., remember ‘left’, click inside rightward box) with the direction of the mouse movement action. The performance difference between compatible and incompatible trials (the ‘compatibility effect’) served as our index of the interplay between WM content and motor behavior. We also varied the proportion of compatible to incompatible trials across blocks of the experiment, so that we could examine the extent to which the compatibility effect is under cognitive control. We then tested how TMS to lateral PFC impacted this control over WM-action interplay.

### Participants

We aimed for a sample of 24 subjects, with three task sessions each, based on a priori sample size estimation. Power analyses were conducted in G*Power 3.1.9.2 (http://www.gpower.hhu.de) for repeated-measures ANOVAs, and we considered a range of relevant previous studies to estimate our expected effect size. We considered a previous study from our group that assessed WM after theta-burst TMS to PFC (Lee & D’Esposito, 2012). This study yielded TMS effect sizes in the range of *f(U)* = 0.51 to 0.78, which corresponds to a medium effect. Studies from other groups that applied theta-burst TMS to frontal or parietal cortex in similar within-subject designs report comparable effect sizes within this range, for both TMS main effects and interactions with task conditions (Heinen et al., 2017; Morgan et al., 2013). We used the mean effect size across these studies (*f(U)* = .62) in power calculations, with the power parameter set to .8, which yielded a total sample size of 24.

We enrolled subjects until we reached this target sample size after exclusions and quality control. Twenty-six subjects (9 male), aged 19-25 (mean: 21.3), were recruited from the Berkeley community, completed informed consent in accordance with the UC Berkeley Institutional Review Board, and were paid $20 per hour for their participation, plus a $20 completion bonus for finishing all sessions. Subjects were screened for MRI and TMS contraindications, and reported no history of neurological, psychiatric, or a significant medical condition. Two subjects were excluded from further sessions — one because orthodontic braces produced severe signal distortion in the MRI, and the other revealed a TMS contraindication after initial scanning. Therefore, 24 subjects remain in the reported analyses.

### Behavioral task stimuli and procedure

The behavioral task (**Fig. 1b**) was nearly identical to that used by Miller et al. (2020; Experiment 1), and the description below recapitulates those methods. The task was programmed and presented using MATLAB (https://www.mathworks.com/), running Psychtoolbox routines (Brainard, 1997; http://psychtoolbox.org/), along with custom scripts to continuously track mouse positions. Stimuli were displayed on a neutral grey background (RGB: [128,128,128]) and viewed from approximately 60 cm. The WM stimuli consisted of directional words (‘up,’ ‘down,’ ‘left,’ or ‘right’) that were displayed in black lettering. Every trial began with a 2 sec intertrial interval (ITI). Then a WM sample word appeared centrally for 1 sec. After a total delay of 5 sec, a WM probe word (selected from the same set as the WM samples) appeared centrally underneath a question mark. The WM task was to make a keyboard button press with the left hand indicating whether the probe word was a match (‘S’ key) or non-match (‘D’ key) to the WM sample. Match and non-match WM probes were equally likely (50% match / 50% non-match) and occurred in random order.

During the WM delay, participants completed a cued hand movement. A central filled colored square (i.e., the cue) was flanked by unfilled square boxes (i.e., the targets) at each of four locations: to the top, bottom, left, and right of center. The central square could be one of four colors (RGB: green = [122,164, 86], pink = [198, 89, 153], orange = [201,109, 68], blue = [119,122, 205]), which were chosen to be maximally distinct, matched on saturation and brightness, and color-blind friendly (http://tools.medialab.sciences-po.fr/iwanthue/). Each color was instructed to cue one of four screen locations: green *= left,* pink *= right,* orange *= up,* blue *= down*. The target boxes were equidistant from the central color cue and from each other (∼3.7° in size, ∼9.3° in distance from center). The motor task was to move the mouse with the right hand and click inside the target box at the location cued by the color. The motor task therefore required a symbolic transformation from color to location, which was meant to engage the goal representation circuitry involved in gating motor behaviors (Oliveira & Ivry, 2008; O’Reilly & Frank, 2006). The motor task epoch ended when a 2 sec response deadline passed.

Together, the sequence of one complete dual-task trial started with a 2 sec ITI, followed by a 1 sec WM sample display, then a 2 sec fixation delay. After this first delay, the motor task display appeared for 2 sec, followed by another fixation delay of 1 sec, and then finally the WM probe display for 2 sec. In the main departure from a previous version of the task (Miller et al., 2020), trial timing was fixed to ensure a consistent task duration, in the effective TMS window, for each participant and session regardless of response time. There were two primary trial types: compatible trials, wherein the meaning of the WM word matched the cued direction of movement, and incompatible trials, wherein the WM word was paired with any of the three non-matching movement cues. The ratio of compatible to incompatible trials was manipulated across a given task block. Blocks contained either 80%, 50%, or 20% compatible trials. In “high compatibility” blocks (80% compatible), the WM sample would usually help the motor task, as it suggested the goal direction for the upcoming movement. In “middle compatibility” blocks (50% compatible), the WM content was equally likely to help or harm to the motor task on any given trial. In “low compatibility” blocks (20% compatible) the WM sample meaning usually differed from the motor task target, and was therefore unhelpful. To minimize variability across participant learning effects, they were explicitly informed about the percentage of compatible trials at the start of each block.

Participants practiced at least 12 trials of the motor task, to learn the color-direction response mapping to criterion, before the first experimental session. In all sessions, participants completed one 5-trial practice block of each dual-task block condition (15 total practice trials, with feedback for motor and WM response accuracy) before completing three 30-trial experimental blocks of each condition (9 blocks total; without feedback). Participants therefore completed 90 trials in each block condition and 135 trials of each compatibility condition across blocks (72 incompatible/18 compatible trials across “low compatibility” blocks, 45 incompatible/45 compatible trials across “middle compatibility” blocks, and 18 incompatible/72 compatible trials across “high compatibility” blocks). This amounted to 270 total trials per subject in each session, and 810 total trials per subject across sessions. The first block in each session was always middle compatibility (50% compatible) and the predictability conditions occurred in random order for the remaining blocks. The difference in motor behavior on compatible vs. incompatible trials—or the ‘compatibility effect’—will serve here as an operational index of the WM influence over ongoing action.

### MRI methods

#### Acquisition

All images were acquired on a research-dedicated Siemens TIM/Trio 3T MRI system in the Henry H. Wheeler, Jr. Brain Imaging Center at the University of California, Berkeley. High-resolution anatomical images were acquired with a T1-weighted MPRAGE sequence (TR: 2300ms; TE: 2.98ms; flip angle: 9°; 160 sagittal slices; 1 × 1 × 1 mm). After the anatomical scan, participants completed one run of closed-eye rest for 9-10 minutes. Functional pulse sequence parameters varied slightly across participants, as some were recruited from other studies within the lab. That is, we already had the necessary scans on file for some participants. The resting functional data are inessential to the main conclusions and are included only in exploratory analyses. Functional data were acquired with a gradient echo T2*-weighted echo-planar imaging (EPI) sequence (*15 subjects* - TR: 2000ms, TE: 23ms, flip angle: 50°, 35 transverse slices, 3 × 3 × 3 mm; *6 subjects* - TR: 2000ms, TE: 32.2ms, flip angle: 45°, 54 transverse slices, 2.5 × 2.5 × 2.5 mm; *3 subjects* - TR: 2000ms, TE: 28ms, flip angle: 78°, 32 transverse slices, 3 x 3 x 3 mm).

#### Analysis

##### TMS target transformation

The target Montreal Neurological Institute (MNI) coordinates were first converted into each participants’ native space for TMS. Each participant’s T1-weighted anatomical MRI was passed through the *SPM12* normalization procedure (https://www.fil.ion.ucl.ac.uk/spm/software/spm12/) to the MNI template space (*normalise* function). The affine transformation mapping was then used to derive the individual native space coordinates that corresponded to the MNI coordinates of the *a priori* target ([-40, 32, 32]).

##### fMRI preprocessing

Exploratory results included in this manuscript come from preprocessing performed using *fMRIPrep* 1.4.0 (Esteban et al., 2019; RRID:SCR_016216), which is based on *Nipype* 1.2.0 (Gorgolewski et al., 2017; RRID:SCR_002502).

##### Resting-state connectivity analysis

For each participant’s fMRI resting-state timeseries, correlation was calculated between the TMS target coordinate (5mm sphere) and all other brain voxels. To calculate correlations, functional data were filtered between 0.01 and 0.1 Hz, detrended and standardized, and the top 5 CompCor components were included as confound regressors (implemented in *Nilearn* python package). A fixed-effects GLM across all participants was then calculated in SPM12 to obtain the voxels with the highest overlap with the target coordinate. The group connectivity map was then clustered in *Nilearn* and the mean values for top clusters were extracted for each participant.

### TMS Methods

#### TMS design

It is difficult to achieve perfect experimental control in TMS research. TMS over different cortical targets can produce vastly different scalp sensations and degrees of participant discomfort (Meteyard & Holmes, 2018). Apparent regional differences in TMS response may further be driven by differences in scalp-to-cortex distance (McConnell et al., 2001) or cortical geometry (Pell et al., 2011) that are unrelated to the function of interest. Moreover, TMS to cortical sites of no experimental interest (which is a common control condition) can propagate throughout networks that are involved in task performance (Jung et al., 2016). These factors can disguise the ground truth and obscure whether stimulation effects are driven by “control” or experimental TMS. Here, we took two measures to achieve experimental control. First, we used two different active TMS protocols (cTBS and iTBS), to the same experimental target site. Unlike sham TMS or control stimulation to a different region — which would both produce noticeably different sensations from the experimental stimulation — cTBS vs. iTBS provide strong active controls for each other. Both protocols deliver the same number of pulses, in bursts of the same frequency, at the same intensity, to the same target region. However, because of differences in the patterning of bursts, the protocols are expected to have opposite effects on cortical excitability (Huang et al., 2005). The different forms of stimulation should therefore produce distinct outcomes in behaviors that depend on the targeted region. Second, we supplemented this active control with a no-TMS condition (where all other procedures were identical), to replicate previous findings and establish the ground truth performance on each task measure.

#### Target selection

Here, we aimed to interrogate the mechanisms that prioritize and deploy WM content for action. The selection and use of WM, or ‘output gating’, has been attributed to a cortico-striatal pathway, specifically to connectivity between the lateral PFC and caudate nucleus (Chatham et al., 2014). We therefore aimed to stimulate this cortico-striatal circuit. We cannot directly target subcortex with TMS, but TMS to the mid-lateral PFC can selectively modulate dopamine synthesis and fMRI activity in the caudate, as compared to more caudal PFC TMS which modulates the putamen (Riddle et al., 2022; Schouwenburg et al., 2012; Strafella et al., 2001, 2003). That is, TMS to distinct frontal cortical targets, with distinct anatomical connectivity profiles, can alter fMRI and neuromodulatory activity in select subcortical nuclei. This sort of ‘sling-shot’ approach has also been reliably applied to cortical temporal and parietal targets to modulate hippocampal activity and memory performance (Tambini et al., 2018; Wang et al., 2014). We chose a left dorsal mid-lateral PFC target region [MNI: −40, 32, 32] with demonstrated anatomical and functional connectivity to striatum, using peak coordinates from prior studies that have been shown to modulate caudate activity in response to TMS (Choi et al., 2012; Petrides et al., 1993; Strafella et al., 2001; **Fig. 1a**). We reasoned that this PFC target would be most likely to engage the circuitry that is relevant to our cognitive functions of interest.

#### TMS equipment and parameters

A Magstim Super Rapid stimulator with two booster modules delivered stimulation via a Magstim Double 70mm Air Film coil. We used the Brainsight 2 frameless stereotactic neuronavigation system (Rogue Research, Montreal, Canada) to localize the TMS targets for each participant and to monitor the coil position throughout each stimulation run. The coil was oriented perpendicular to the underlying gyral/sulcal anatomy, and the coil handle was positioned posterior to the coil head.

To individually calibrate the stimulation intensity for each participant, we assessed motor threshold (MT) using electromyographic recording of the dorsal interosseus muscle of the right hand. Single pulses were delivered over the hand representation area of left primary motor cortex, which was first approximated as 5cm lateral from the vertex. Then a search procedure, moving along a grid around that spot, identified the location where stimulation produced the greatest motor-evoked potential (MEP) when the muscle was relaxed. This was defined as the motor “hot spot.” The hot-spot was then used to measure active motor threshold: the minimum single-pulse stimulation intensity required to produce 5 out of 10 MEPs, with peak-to-peak amplitude of at least 200μV, while the participant maintained a voluntarily muscle contraction of ∼20%. A raw EMG display provided the participant with continuous visual feedback to facilitate consistent force of muscle contraction. Active MT ranged from 36-65 percent of maximum stimulator output (mean: 45.2) across this sample.

Following published procedures and safety guidelines (Oberman et al., 2011), the stimulation intensity was set at 80% of each participant’s active motor threshold. However, we had difficulty reaching a stable threshold for two participants. At high stimulation intensities we observed occasional large MEPs, but few twitches in the contralateral finger and few MEPs within the specified thresholding range. In order to preserve participant comfort and safety, we capped TBS intensity at 44% MSO. Therefore, stimulation intensities ranged from 29-44% in this sample.

The two experimental TMS sessions delivered an offline theta-burst TMS protocol (Huang et al., 2005), which can result in up to 60 min of impact on cortical excitability of the targeted region (Gamboa et al., 2010; Gentner et al., 2008; Grossheinrich et al., 2009). Theta-burst stimulation (TBS) comprises trains of 3 pulses at 50 Hz, delivered once every 200 msec (i.e., at theta frequency). TBS is delivered in two primary patterns: continuous TBS (cTBS) and intermittent TBS (iTBS). cTBS involves a continuous train of TBS for a total duration of 40 sec, whereas iTBS involves 2 sec trains of TBS repeated every 10 seconds (i.e., with an 8 sec break between trains), for a total of 190 seconds. Both cTBS and iTBS protocols deliver a total of 600 pulses. Whereas cTBS has been found to produce transient decreases in cortical excitability, iTBS is instead associated with transient increases in excitability. Moreover, these protocols, delivered to the same frontal region, have shown distinct effects on striatal dopamine release (Aceves-Serrano et al., 2022). While individuals are variable in their degree and direction of TMS effects (Cantone et al., 2019; Corp et al., 2020; Hamada et al., 2013; Tik et al., 2023), cTBS and iTBS are generally expected to have distinct effects on cortical excitability when applied over the same cortical region. This experimental design should therefore produce distinct behavioral effects of the stimulation conditions, if the behavior depends on the lateral PFC target.

For each TMS session, participants first completed 15 trials of behavioral task practice immediately before stimulation, then underwent the TBS procedure. One experimenter delivered the stimulation while another monitored the participant for comfort and safety. We observed facial twitching during stimulation in 5 (out of 24) participants, but no participants reported pain and we observed no other adverse effects. When the TBS protocol ended, participants rested for 5 minutes before completing the behavioral task, which lasted 48 minutes.

### Analysis strategy

#### Movement trajectory measures

We continuously tracked the mouse position across each trial of the motor task (**Fig. 2**). To evaluate the precision of motor trajectories, we defined a circular boundary around the central start position, with a radius of ¼ the distance to the target (**Fig. 2b**), and we measured the angle at which the cursor crossed that boundary. We calculated the deviation from the target direction, and **Fig. 2a** shows the distribution of these angular errors for all compatible and incompatible trials. To quantify trajectories that initially curved away from the target location, movements were considered precise if they first crossed the circular boundary within 45° of the correct response axis, but were classified as course adjustments if they crossed the boundary at a wider angle than 45° before terminating at the correct target (**Fig. 2b**). All cursor trajectories were rotated to a common axis for comparison.

**Figure 2.**
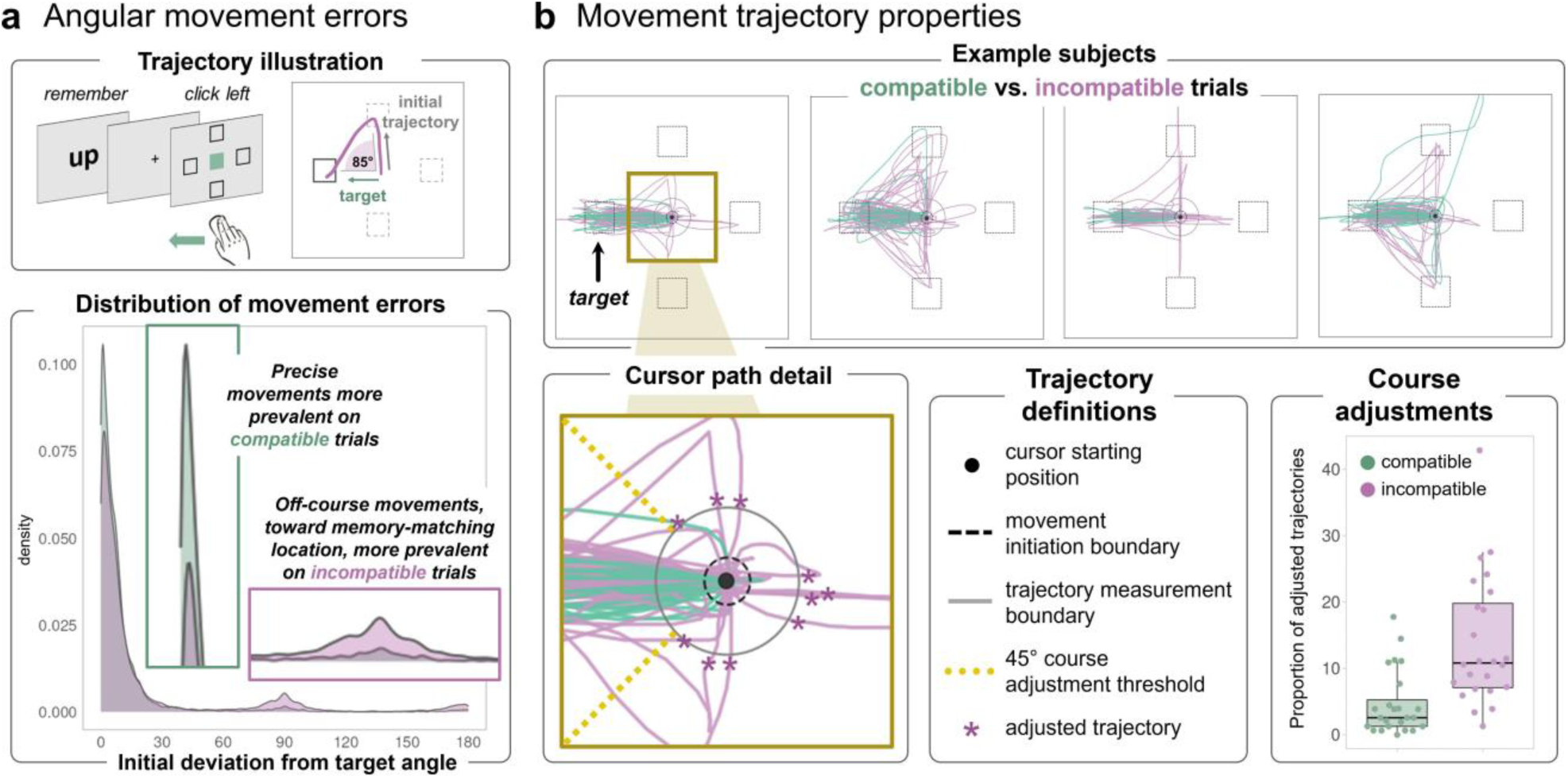
Dissecting discrete properties of movement behavior. (a) We continuously tracked mouse trajectories after the onset of the color movement cue. We calculated the initial deviation away from the target location, and display the frequency distribution of angular errors, for compatible and incompatible trials. (b) Movement trajectories for compatible (green) and incompatible trials (purple) from example subjects, in middle compatibility blocks (rotated to a common target direction). Detail illustrates boundaries for determining movement initiation, measuring angular error, and categorizing trials as course adjustments. Course adjustments were more frequent on incompatible trials (shown for middle compatibility blocks, when compatible and incompatible trials occurred equally often).

#### Movement speed measures

Movement initiation—also sometimes referred to as ‘reaction time’—was defined as the time from the onset of the color cue until the cursor first crossed a radius of 30 pixels from the starting position (**Fig. 2b**). Movement duration—also sometimes referred to as ‘movement time’—was defined as the amount of time after movement initiation until a click was made in any target box.

#### QA

For all experiments and measures, we excluded trials that were missing a WM probe response. For movement speed analyses, we excluded outlier trials when a measurement was greater than 3 standard deviations away from the participants’ mean, or if the motor task response landed at the incorrect target location.

#### Statistics

We specified two sets of mixed-effects models to test the TMS influence on WM-motor compatibility effects and block-level control. First, we used a set of ‘three-target’ models that included the *no TMS* condition, so that both experimental TMS conditions could be compared to ground truth behavior. We constructed linear models with the response time variable of interest (e.g., movement initiation, movement duration) as the outcome measure, and we coded predictor variables as PFC *TMS protocol* (treatment coded with the *no TMS* condition as the relative baseline), *trial compatibility* (compatible vs. incompatible), *block proportion* (continuous measure of compatibility proportion for each block, mean-centered), and *block number* (mean-centered), including interactions. For WM or movement accuracy, we used a logistic mixed model with the same predictor variable coding. All mixed linear models were constructed with subject as a random effect (random-intercepts), and built and tested via the *lme4* (https://github.com/lme4/lme4/) and *lmerTest* (https://rdrr.io/cran/lmerTest/) R packages, ported in Python via ryp2 (https://rpy2.github.io/).

We then constructed ‘two-target’ models with the same structure, but including only the active TMS conditions. These were designed to directly compare cTBS vs. iTBS and account for non-specific effects of stimulation. Both two- and three-target models reveal generally consistent outcomes, so are included to convey a full picture of the conditions that drive significant effects. In visualizations we include the *no TMS* baseline alongside the experimental TMS conditions, but conclusions are based on findings that differ between the two experimental TMS conditions (cTBS and iTBS in the two-target model).

We also specified a final set of models to test the TMS influence on trial compatibility sequence effects – that is, the influence of the preceding trial condition on current trial compatibility effects. We selected only the neutral/middle compatibility block conditions (block compatibility = 50%) from each session and coded an additional variable for *previous trial compatibility* (compatible vs. incompatible). In these models, PFC *TMS protocol* (either 3- or 2-way coded as above), *current trial compatibility*, and *previous trial compatibility* were included as explanatory variables.

For the fixed effects of each model, we report a parameter estimate value, estimated confidence interval (t-tests from *lmerTest* using Satterthwaite’s approximation (Kuznetsova et al., 2017)), and *p*-value. Full model specifications and fitting results are listed in *Extended Data Tables*.

## Results

### Baseline condition replicates previous findings

We first aimed to replicate our previous behavioral findings, so we applied the prior study analysis strategy (Miller et al., 2020) to the data from the *no TMS* condition here. This was meant to ensure that basic WM-motor effects are reliable across studies and cohorts. For all replication analyses, we conducted 2 (*trial compatibility*: compatible vs. incompatible) × 3 (*block proportion compatible*: high vs. middle vs. low) repeated measures ANOVAs, to mirror the previous study.

Motor decision accuracy (i.e., % correct of click location) was slightly worse on incompatible (98%) vs. compatible trials (99%), *F*(1,23) = 22.83, *p* < .001, η_p_^2^ = .18, but there was neither a main effect of *block proportion*, *F*(2,46) = 1.4, *p* = .25, η_p_^2^ = .02, nor an interaction between factors, *F*(2,46) = .11, *p* = .89, η_p_^2^ = .001. When the meaning of the WM content was incompatible with motor goals, participants chose the wrong target location more often. Throughout the manuscript, we refer to this difference between compatible and incompatible conditions as the ‘compatibility effect’.

Movement landing positions were overall highly accurate (98% correct), but WM content may bias the shape of movement trajectories. For instance, incompatible trials may result in curved paths that are skewed toward the direction that matches the meaning of the WM sample. Indeed, the proportion of movement course adjustments was ∼10% greater on incompatible versus compatible trials across all block conditions, *F*(1,23) = 33.71, *p* < .001, η_p_^2^ = .41 (**Fig. 2b**). There was no effect of *block proportion* condition (*p* = .2), but there was an interaction between *trial* and *block* factors *F*(2,46) = 7.76, *p* = .001, η_p_^2^ = .033. Action slips on incompatible trials were especially likely in high compatibility blocks when WM content was most likely to match movement goals.

WM content or its relevance to the motor task could influence multiple ongoing processes. We therefore examined distinct *movement initiation* and *duration* measures geared to index motor decision-making vs. movement execution speed, respectively. A main effect of *trial compatibility*, *F*(1,23) = 45.26, *p* < 001, η_p_^2^ = .35, indicated that movements were *initiated* more slowly when the cued movement was incompatible with the WM sample meaning (67 ms difference). A main effect of *block proportion*, *F*(2,46) = 17.16, *p* < 001, η_p_^2^ = .15, indicated that movements were also initiated more quickly overall in blocks when WM content was more likely to help motor performance. Moreover, there was an interaction between *trial compatibility* and *block proportion* factors, *F*(2,46) = 3.37, *p* = .043, η_p_^2^ = .016, whereby the compatibility effect was amplified in blocks with more compatible trials. Compatible movement initiation was especially speeded when those trials were most likely (**Fig. 3a**).

**Figure 3.**
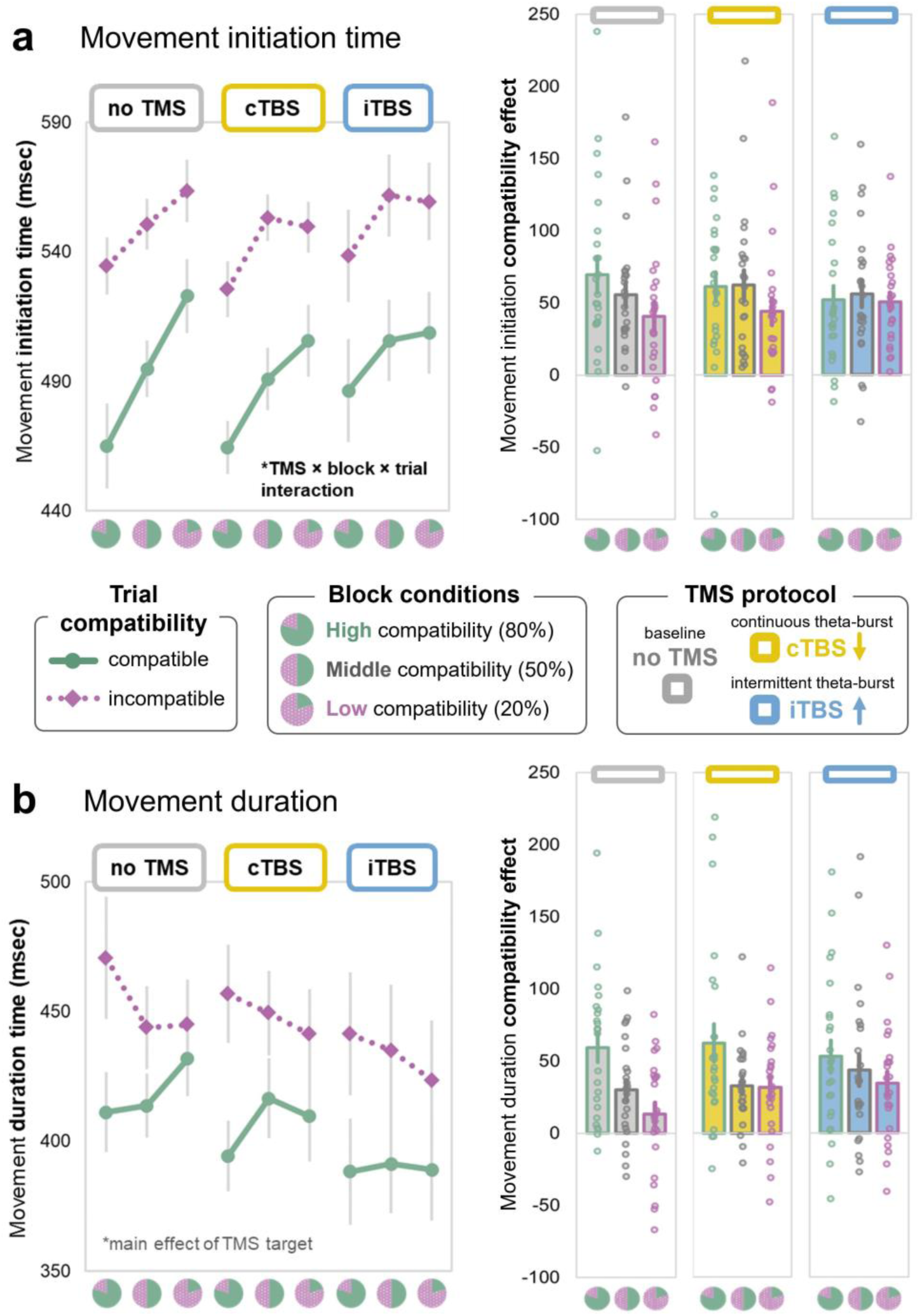
TMS effects on block-level cognitive control. Left panels illustrate raw group means for each TMS protocol, block proportion, and trial compatibility condition. Right panels illustrate compatibility effect difference scores (incompatible – compatible) for movement measures. (a) Movement initiation time. (b) Movement duration. Points reflect each individual subject average for that condition, and error bars reflect SEM.

*Movement duration* also showed a main effect of *trial compatibility*, *F*(1,23) = 26.95, *p* < .001, η_p_^2^ = .22. Movements took longer on incompatible trials (38 msec difference).

However, unlike movement *initiation* — which was overall speeded by a higher frequency of compatible trials — movement *duration* was slowed instead. There was an interaction between *trial* and *block* factors, *F*(2,46) = 12.34, *p* < .001, η_p_^2^ = .067, where the compatibility effect was amplified in blocks with more compatible trials. Incompatible movement duration was especially slowed when those trials were least likely (**Fig. 3b**).

#### Replication Summary

These results show a near-identical replication of the response pattern in our previous behavioral study, wherein motor benefits of compatible WM content manifested in accelerated decision processes, while costs of incompatible content manifested in more imprecise and time-consuming actions. Therefore, the *no TMS* condition represents a stable baseline against which to compare behavior after experimental TMS.

### TMS influences block-level (proactive) cognitive control

The replication analysis showed a systematic pattern of trial and block condition effects, whereby the magnitude of compatibility effect was modulated by the likelihood of compatible/incompatible trials: the greater the proportion of incompatible trials, the less they impacted movement speed and accuracy. This pattern may be considered to reflect sustained, anticipatory control over WM-action conflict at the level of the task block context (i.e., proactive control; Braver, 2012). This pattern was altered after TMS to mid-lateral PFC.

#### Movement choice and trajectory

In the original study, as well as our replication in the *no TMS* condition, movement target choice accuracy was high, but displayed a small compatibility effect. Here, TMS also produced an interaction with trial compatibility and time across the task (i.e., block number, which was included in the model to capture that neural effects of TMS are expected to decay over time). PFC-targeted cTBS modulated the magnitude of compatibility effect across the course of the task relative to a *no TMS* comparison (ß = 0.005, *p* = 0.044), and in a direct iTBS vs. cTBS comparison (ß = −0.005, *p* = 0.049; see *Extd. Data Tables*). The compatibility effect (accuracy for compatible vs. incompatible trials) was magnified after cTBS, especially in earlier task blocks. This cTBS effect decreased over time, and there was no TMS interaction with block proportion compatibility. TMS condition (either relative to *no TMS* or cTBS vs. iTBS) also had no impact on the precision or course of correctly-chosen movements [*p-values* > 0.2, see *Extd. Data*].

#### Movement speed

In the original study, and in the *no TMS* replication, movement initiation was more conservative in blocks when incompatible trials were more likely. Even compatible trial movements started slower in those blocks, suggesting a temporally-extended tendency for response caution. Conversely, movement initiation was overall speeded in blocks when incompatible trials were less likely, and this effect was magnified for compatible trials, suggesting a lower movement decision threshold when WM and movement goals tended to align. However, that block level (proactive) modulation was attenuated after TMS (**Fig. 3a**). Movement initiation speed was impacted by a combination of TMS target, block, and trial compatibility conditions (see *Extd. Data Tables* for all tests), including a *TMS* x *trial compatibility* x *block proportion x block number* interaction in both models (ß = −26.4 [CI: −44.8, −8], *p* = 0.005; 2-target model), as well as a main effect of *TMS* protocol (ß = 13.8 [CI: −6.4, 21.2], *p* < 0.001;). More specifically, after iTBS, there was no relative slowing when compatible trials were least likely, and there was less benefit to compatible movement initiation when compatible trials were most likely. Movement *duration*, however, showed no trial- or block-type specific TMS effects. Instead, a graded effect of *TMS* protocol indicated that movements were executed faster overall after iTBS (which is meant to be excitatory; **Fig. 3b**) relative to both cTBS (ß = −18.6 [CI: −26.7, −10.5], *p* < 0.001) and the *no TMS* baseline (ß = −28.4 [CI: −36.6, −20.2], *p* < 0.001).

#### WM recognition

WM probe accuracy was high overall, averaging 95-99% across all TMS and task conditions, and unharmed by either stimulation condition. Both probe accuracy and RT were worse on incompatible trials, but no other block or TMS effects emerged in the two-target model (*p-values* > 0.2, see *Extd. Data*). In fact, the three-target model revealed that WM probe recognition was slightly more accurate on incompatible trials after iTBS (compared to *no TMS* baseline: ß = 0.02 [CI: 0.01, 0.03], *p* = 0.003). Therefore, cognitive control in response to block-level trial proportions seems to affect the intervening movement, but not memory performance. Moreover, TMS seems to modulate the effect of WM on intervening movement behavior, without harming the memory itself.

##### Interim Summary

Compatibility effects in movement choice accuracy were amplified by cTBS to mid-lateral PFC, while trajectory precision was left unaffected. TMS also modulated block-level (i.e., proactive) cognitive control over WM biases in movement speed. iTBS impacted condition-specific control over movement initiation, while movement duration showed a non-specific acceleration. iTBS seemed to eliminate context-sensitive cognitive control settings that ignite or check movement initiation depending on the relationship between internal content and action goals. These condition-specific effects on movement choice and initiation time – but not trajectory precision or movement duration – suggest that TMS modulated decision-making processes, but not movement execution. WM probe accuracy remained high in all conditions, indicating that TMS effects on conflict control are unlikely due to impacts on the WM representation. Instead, the stimulated PFC region seems to uniquely modulate the interplay between WM and actions, depending on their likelihood of overlap.

### TMS influences trial-to-trial (reactive) cognitive control

As shown above, TMS to lateral PFC modulates sustained cognitive control over WM-motor interactions (as operationalized by block-level proportion compatibility effects). In our previous study, we also delivered trial-level manipulations of item priority, to examine control processes on a more immediate time scale. We found that temporally-extended, higher-order task goals exerted distinct effects vs. phasic, item-level attentional modulation (Miller et al., 2020). We, therefore, reason that PFC may make distinct contributions to block-vs. trial-level conflict control processes here (Braver, 2012). ‘Congruency sequence effects’ describe a phenomenon observed in conflict tasks like the Stroop or flanker, whereby the congruency effect on a given trial interacts with the congruency condition of the preceding trial (Egner, 2007; Gratton et al., 1992). Conflict effects tend to be magnified after a congruent trial, while they are dampened after an incongruent trial – presumably because the conflict triggers a reactive upregulation in cognitive control. Here, we extend this framework to illuminate the principles of interplay between WM content and motor behavior. We examine the effects of short-term context (i.e., the preceding trial) on WM-motor compatibility effects, and test the contribution of lateral PFC to such a reactive, trial-by-trial form of WM-action control. For these analyses, we focus only on middle compatibility task blocks (50%), wherein compatible and incompatible trials occurred equally often.

#### Movement choice and trajectory

In the *no TMS* condition, movement accuracy displayed a typical descriptive pattern of congruency sequence effects (**Fig. 4a**), but many participants achieved 100% accuracy (and therefore showed no conditional effects). There were no significant effects of previous trial compatibility or TMS condition for either choice accuracy [*p-values* > 0.2], or course trajectory adjustments [*p-values* > 0.2, see *Extd. Data*].

**Figure 4.**
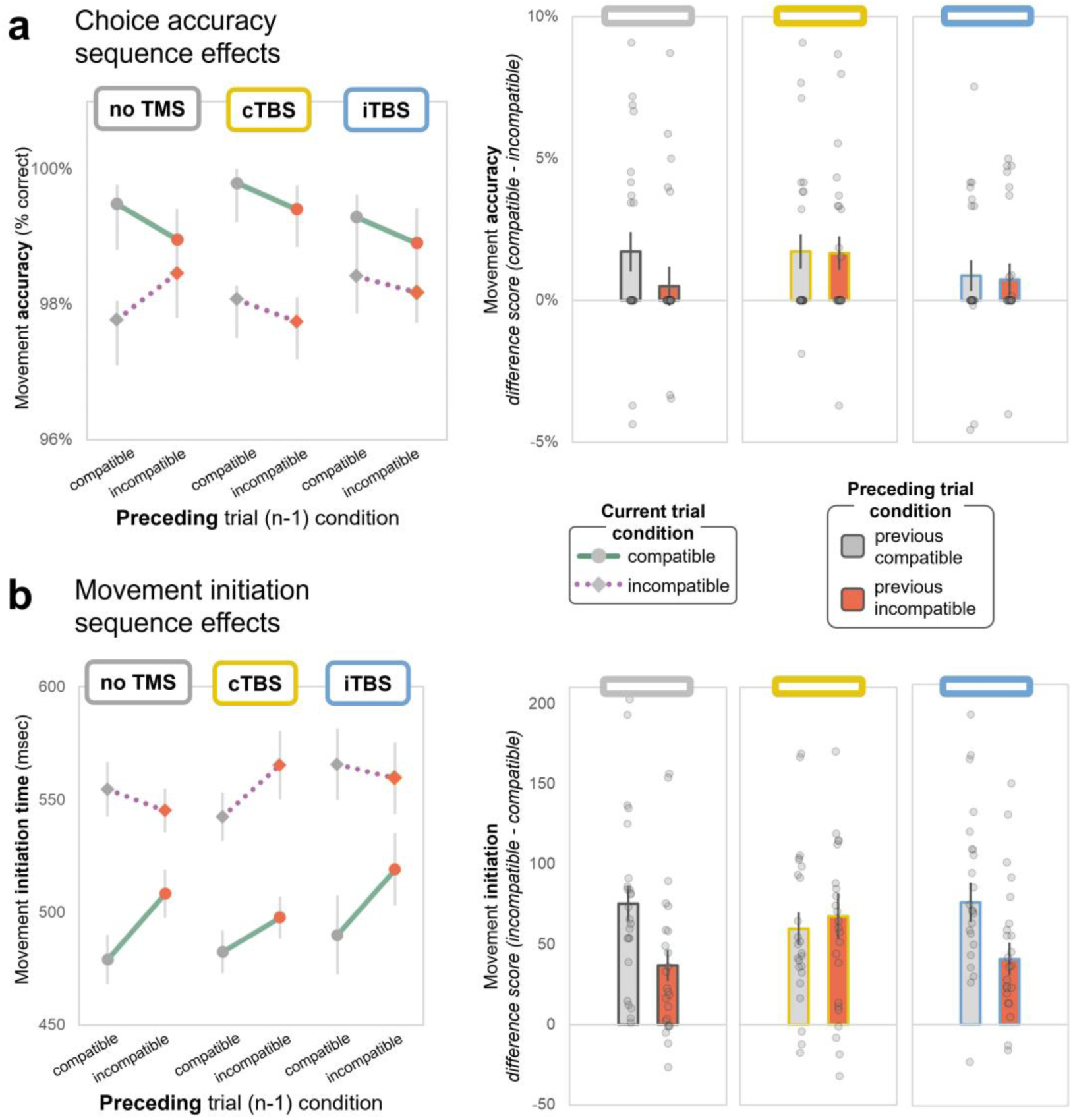
TMS effects on trial-by-trial compatibility sequence effects (CSE). CSE analyses were conducted only for middle compatibility blocks, wherein both trial types are equally likely. (a) Movement choice accuracy group means for each TMS condition, as a function of both current and previous trial compatibility (left). Different current trial compatibility conditions are displayed with separate lines, while previous (n-1) trial compatibility conditions are displayed with grey vs. red markers. Compatibility effect difference scores (compatible – incompatible) for each TMS and previous trial condition are shown on the right. (b) Movement initiation group means for each TMS condition, as a function of both current and previous trial compatibility (left). Compatibility effect differences scores (incompatible – compatible) for each TMS and previous trial condition (right). Points reflect each individual subject average for that condition, and error bars reflect SEM.

#### Movement speed

In the *no TMS* condition, the sequence effects that are characteristic of conflict scenarios (Egner, 2007) emerged here in trial-to-trial control over WM-motor interactions. The movement initiation compatibility effect was smaller following incompatible (vs. compatible) trials. Immediately after having experienced incompatible trial conflict, compatible movements were initiated more slowly (relative to when the preceding trial was compatible) while incompatible movements were initiated more quickly (*current trial* x *previous trial compatibility* interaction; ß = −41.6 [CI: −64.4, − 18.9], *p* < 0.001; **Fig. 4b**). This suggests that conflict between WM content and motor goals triggered a control adjustment to dampen the influence of WM content (whether helpful or harmful) on subsequent trials.

This distinctive marker of adaptive control was eliminated, and descriptively reversed, after cTBS (*TMS* x *current trial* x *previous trial compatibility* interaction; three-target model: ß = −43.7 [CI: 11.7, 75.8], *p* = 0.008; two target model: ß = −39.3 [CI: - 71.3, −7.2], *p* = 0.016). Movement initiation became overall slower after incompatible trials, and the magnitude of compatibility effect was equivalent whether the previous trial was compatible or incompatible (**Fig. 4b**). That is, rather than a reactive improvement to incompatible responding after conflict (as would be expected under typical function), movement initiation was delayed instead. This suggests that PFC cTBS interferes with adaptive control that would otherwise enhance or dampen the influence of WM content over movements, in response to compatible vs. incompatible trials respectively. Under cTBS, incompatible movement initiation was especially hindered after incompatible trials.

Movement duration displayed no such compatibility sequence effects and, like the block-level control effects, PFC stimulation had no impact on trial-by-trial control over movement duration [*p-values* > 0.2, see *Extd. Data*].

##### Interim Summary

Conflict between dual-task WM content and action goals displayed characteristic congruency sequence effects (in movement initiation time) and TMS to mid-lateral PFC modulated this trial-by-trial control. cTBS, which is meant to be inhibitory, eliminated the sensitivity to short-term control demands which is typically expressed as a smaller compatibility effect after conflict trials. Instead, cTBS resulted in overall slowing after incompatible trials, and a descriptively larger compatibility effect. Choice accuracy effects failed to meet statistical thresholds, but descriptively mirrored the pattern in movement initiation. Once again, TMS effects were limited to measures that index decision processes. However, this reactive trial-by-trial control effect was selectively modulated by cTBS, whereas the block-level control effects (in the previous section) were modulated more by iTBS instead.

### Exploratory analyses of individual variability

This study was not powered to formally examine individual differences in behavior, but it is now understood that TMS effects can vary widely with properties of the individual, as well as their brain state (Corp et al., 2020; Silvanto et al., 2008; Silvanto & Pascual-Leone, 2008). Connectivity between the targeted region and other task-relevant areas may be an important explanatory factor in stimulation effects (Castrillon et al., 2020). We therefore report some exploratory brain-behavior correlations to guide future inquiry.

#### Functional connectivity with the PFC target

TMS effects are not confined to the targeted region, and instead can propagate throughout a network (Eldaief et al., 2023; Hebscher & Voss, 2022). In fact, we hoped to capitalize on this property to engage the cortico-striatal circuitry that is thought to support the behavior we test here (Chatham et al., 2014; Strafella et al., 2001). Moreover, cTBS and iTBS may be mediated by different connections, and induce distinct physiological effects (Aceves-Serrano et al., 2022; Cocchi et al., 2015). Therefore, we reasoned that the TMS effects observed here may further correlate with the strength of individual functional connectivity between the targeted left lateral PFC region and other putative control regions. Different connections may, moreover, be important for different aspects of performance, namely proactive vs. reactive forms of cognitive control. First, we identified regions that were most strongly correlated with the TMS target site during resting-state fMRI (**Fig. 5a**). From that map, we then extracted individual connectivity coefficients with regions-of-interest that are typically implicated in WM and cognitive control – in this case, a parietal cluster and the dorsal medial PFC (dmPFC, encompassing anterior cingulate cortex). We then correlated those connectivity metrics with measures of block- and trial-level cognitive control, which were modulated by iTBS and cTBS respectively (**Fig. 5b**). We focused on compatibility effects in movement initiation, since that measure was most consistently modulated by TMS across analyses. We found that current trial compatibility effects after cTBS (but not iTBS) correlated positively with dmPFC connectivity (Pearson’s *r* = .45, *p* = .027), but not with parietal (*r* = −.26, *p* = .22). That is, stronger connectivity between the TMS target and dmPFC was associated with larger individual TMS-induced compatibility effects (i.e., a marker of susceptibility to conflict). We also found that previous trial compatibility effects after iTBS (but not cTBS) correlated negatively with parietal connectivity (*r* = −.52, *p* = .01), but not with dmPFC (*r* = −.05, *p* = .8). That is, stronger connectivity between the TMS target and the left parietal cluster was associated with smaller TMS-induced compatibility effects after incompatible trials (i.e., a marker of adaptive control). These are meant as only exploratory correlations, but they corroborate the idea that cTBS and iTBS impacts on block- and trial-level control measures may arise through distinct prefrontal circuits.

**Figure 5.**
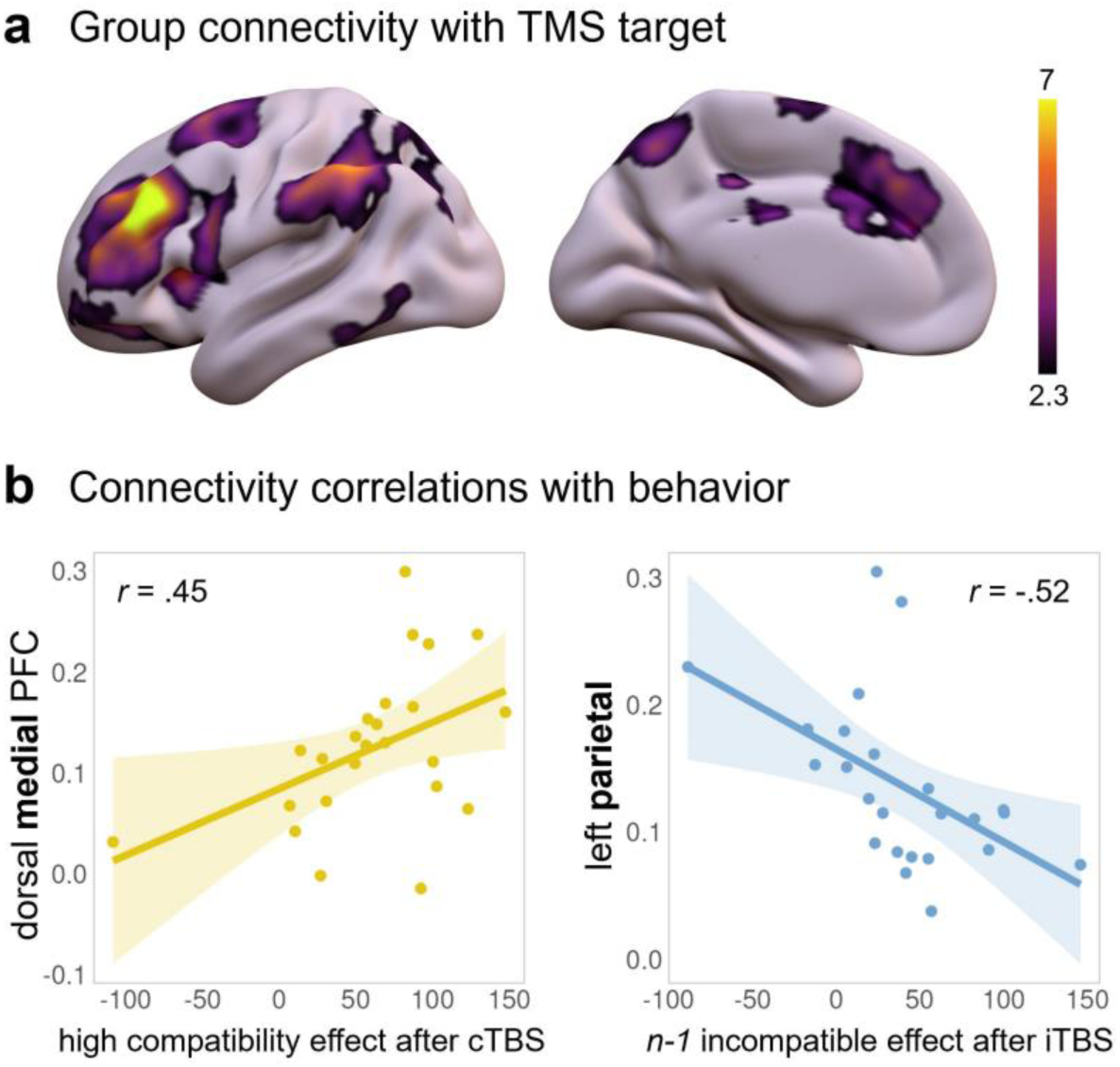
Resting state connectivity with TMS target site. (a) Group-level functional connectivity from the left lateral PFC seed region that was targeted with TMS. (b) Correlations between individual connectivity coefficients and behavioral metrics. Stronger connectivity with the dorsal medial PFC was associated with larger individual compatibility effects, in high compatibility blocks, after cTBS. Stronger connectivity with a left parietal cluster was associated with smaller compatibility effects, following incompatible trials, after iTBS.

## Discussion

Here, we confirmed that verbal WM content can skew the speed and trajectory of manual actions, indicating that WM may unintentionally shape ongoing movements. However, this bias can be minimized when conflict between goals is predictable, so it is also under a degree of strategic control. We then tested the role of left lateral PFC in modulating this controlled interplay between WM and ongoing behavior. We perturbed the region using two distinct TMS protocols (which should have different effects on cortical excitability), and we examined the resultant effects on the type and magnitude of WM-motor interactions. PFC TMS modulated indices of control over movement choice accuracy and movement initiation time, but not movement trajectories or duration. Moreover, iTBS dampened sensitivity to temporally-extended, block-level task contingencies (i.e., proactive control), and cTBS dampened immediate, flexible adaptation to trial-by-trial conflict (i.e., reactive control). Therefore, the same region of left lateral PFC may implement both proactive and reactive forms of control over the interplay between WM and actions, but may do so through different routes.

### Adaptive control over conflict between WM and actions

The behavior observed here resembles the effects of cognitive control in tasks like the Stroop, flanker, or Simon, wherein prepotent stimulus and response tendencies may be in conflict (Braem et al., 2019). The difference here is that the conflicting information is not currently being perceived, but is maintained with WM instead. These findings reaffirm a tight linkage between WM and motor behavior (van Ede, 2020), but now shed light on the processes by which that linkage is controlled when multiple task goals vie for priority.

We find that WM content can produce conflict in motor behavior much like perceived stimuli do, and that interplay appears to follow similar control principles to visual stimulus-response conflict. As in, WM-motor interactions are subject to both proactive control in response to block-level compatibility contingencies, and reactive control in response to recently experienced conflict. In both cases, however, WM performance is unaffected by the control processes that regulate the motor bias. For instance, WM probe accuracy is unaffected by block-level proportion compatibility, suggesting that cognitive control does not suppress interfering WM content per se. Control may instead act to differentiate or integrate WM and motor task sets depending on their likelihood of overlap, while preserving WM integrity (Cole et al., 2017). For instance, when WM and motor goals are likely to be compatible, the gate between them might be open, promoting crosstalk, which would further facilitate compatible movements. However, that crosstalk would also yield inadvertent spillover of WM content into incompatible actions (cf. Hillman et al., 2024). In daily life, this might manifest as inappropriate keystrokes during typing, either because of a lax control state or a high degree of overlap between our ongoing thoughts and typing goals.

### Mid-lateral PFC contributions to memory, decision, and action processes

PFC TMS modulated cognitive control over WM-motor interactions. TMS left WM content recognition unharmed, and resulted in a general speeding of action execution and WM probe responses. However, block- and trial-type specific TMS effects emerged in choice and movement initiation time measures that likely index decision processes. That is, rather than affecting memory or the movement itself, PFC TMS selectively influenced measures of cognitive control that facilitate or limit the sway of WM content on motor decisions. Lateral PFC, therefore, likely mediates the WM-motor transformation instead of underpinning either WM or movement components alone. This is consistent with findings that lateral PFC lesions or transient perturbation often leave essential WM maintenance unharmed, but impair functions that involve manipulating or otherwise controlling WM content (D’Esposito et al., 2006; D’Esposito & Postle, 1999; Mackey et al., 2016). Here, PFC may enable prioritized WM content to bias evidence accumulation for motor decision-making, whereas that decision-making is postponed when WM content is less relevant (Shushruth et al., 2022). We targeted only a mid-lateral PFC node associated with WM output gating. However, targeting a more caudal fronto-putamen circuit might be expected to influence control over movement duration or trajectory instead (Landau et al., 2009).

### Mid-lateral PFC contributions to sustained vs. responsive adaptive control

The targeted PFC region is also considered a processing nexus, with diverse response properties and projections (Chaudhuri et al., 2015; Dang et al., 2021; Duncan, 2001; Miller & Fusi, 2013; Wang et al., 2023; Wasmuht et al., 2018), that coordinates between abstract task goals and immediate concrete action plans (Nee & D’Esposito, 2016). By changing the excitability of the target region, TMS may shift the timescale of PFC processing or alter connectivity with network regions that support more or less temporally abstracted levels of control (Badre & Nee, 2018; Soltani et al., 2021; Soltani & Koechlin, 2022). iTBS, which is meant to be excitatory, interfered with anticipatory control that happens over a longer timescale — eliminating both costs and benefits that depend on higher-order task contingencies. cTBS, which is meant to be inhibitory, instead dampened responsive control that happens over a shorter timescale — eliminating reactive conflict adaptation and subsequent control benefits. WM-related PFC function is characterized by a trade-off between representational stability (which may share mechanisms with proactive control) vs. flexible updating (which may have more in common with reactive control). Excitatory or inhibitory TMS may push the scales in favor of a more stable or flexible prefrontal regime. When iTBS (presumably) increases PFC excitability, it reduces protracted task stability, but when cTBS (presumably) decreases PFC excitability, it reduces rapid task flexibility. This insight could help reconcile competing takes on PFC coding schemes and activity dynamics (e.g., Adam et al., 2023; Christophel et al., 2017; Constantinidis et al., 2018; Leavitt et al., 2017; Lundqvist et al., 2018; Stokes, 2015; Stroud et al., 2024) in that they may, in part, be a product of malleability in PFC control mode.

### TBS effects are individually variable

Here, we aimed to excite or inhibit a PFC-striatal circuit, but the downstream effects of our stimulation are unknown. The directional effects of cTBS and iTBS can be widely variable across people, and we cannot definitively link our outcomes to cortical facilitation or inhibition. Nonetheless, the two protocols did evince distinct behavioral outcomes in this sample, suggesting that they promoted distinct effects on neural activity as intended. The two TMS protocols may have their influence by targeting distinct prefrontal cell types, propagating through different connected regions, or provoking distinct compensatory responses. Indeed, our exploratory brain-behavior correlations indicate that behavioral metrics after cTBS and iTBS were associated with unique functional connectivity profiles across individuals. Stronger dmPFC connectivity was associated with heightened individual susceptibility to conflict after cTBS, while stronger parietal connectivity was associated with dampened individual conflict adaptation effects after iTBS. dmPFC is often implicated in conflict detection and resolution (e.g., in Stroop and flanker tasks; Botvinick et al., 2004), while posterior parietal cortex is implicated in maintaining recent trial history (e.g., Akrami et al., 2018). Stronger connectivity with these control network regions might therefore facilitate stronger relative TMS propagation or compensation depending on the current control demands. More extensive examination of the PFC network pathways for different timescales of WM-motor control, and tests of individual variability across a larger sample should be fruitful areas for future inquiry.

### Conclusions and implications

Here, we tested a simple WM-motor interaction and found that the semantic meaning of WM content biased intervening hand movements. WM can, therefore, hold sway over ongoing motor behavior. Under typical PFC function, this interplay between WM and actions is thermostatically regulated by changes in recent trial conflict history as well as the conflict statistics of the overarching task context. Lateral PFC mediates both immediate (trial history) and overarching (block proportion compatibility) forms of control to adaptively promote or prevent WM content from guiding actions. However, perturbing normal PFC function may alter the balance between proactive and reactive control. This suggests that the same PFC region supports distinct forms of control, but through different network pathways, and changing PFC excitability can change the timescale of control.

## Data and code availability

Data and analysis code will be publicly shared in an OSF repository (https://osf.io/jcytw/) upon publication.

## Acknowledgements

We thank Rich Ivry for collaboration and guidance on task design and movement trajectory analysis approaches.

## Extended Data Tables

### Three-target models (no TMS vs. cTBS vs. iTBS)

**Table 1a.** Movement choice accuracy.

| *Predictors* | *Estimates* | *CI* | *p* |
| --- | --- | --- | --- |
| (Intercept) | 1.00 | 0.99 – 1.00 | <**0.001** |
| target [PFC_cTBS] | -0.00 | -0.01 – 0.00 | 0.292 |
| target [PFC_iTBS] | -0.00 | -0.01 – 0.00 | 0.334 |
| TrialType2 | -0.01 | -0.02 – -0.01 | <**0.001** |
| BlockType_continuous_centered | 0.00 | -0.01 – 0.02 | 0.625 |
| block_centered | 0.00 | -0.00 – 0.00 | 0.681 |
| target [PFC_cTBS] : TrialType2 | 0.00 | -0.01 – 0.01 | 0.776 |
| target [PFC_iTBS] : TrialType2 | 0.00 | -0.00 – 0.01 | 0.531 |
| target [PFC_cTBS] * BlockType_continuous_centered | 0.00 | -0.02 – 0.02 | 0.992 |
| target [PFC_iTBS] * BlockType_continuous_centered | 0.01 | -0.01 – 0.03 | 0.389 |
| TrialType2 * BlockType_continuous_centered | -0.01 | -0.03 – 0.02 | 0.624 |
| target [PFC_cTBS] * block_centered | -0.00 | -0.00 – 0.00 | 0.078 |
| target [PFC_iTBS] * block_centered | -0.00 | -0.00 – 0.00 | 0.967 |
| TrialType2 * block_centered | -0.00 | -0.00 – 0.00 | 0.971 |
| BlockType_continuous_centered * block_centered | -0.00 | -0.01 – 0.01 | 0.865 |
| (target [PFC_cTBS] * TrialType2) * BlockType_continuous_centered | -0.02 | -0.05 – 0.01 | 0.149 |
| (target [PFC_iTBS] * TrialType2) * BlockType_continuous_centered | -0.02 | -0.05 – 0.01 | 0.244 |
| (target [PFC_cTBS] * TrialType2) * block_centered | 0.00 | 0.00 – 0.01 | **0.044** |
| (target [PFC_iTBS] * TrialType2) * block_centered | 0.00 | -0.00 – 0.00 | 0.979 |
| (target [PFC_cTBS] * BlockType_continuous_centered) * block_centered | 0.00 | -0.01 – 0.01 | 0.496 |
| (target [PFC_iTBS] * BlockType_continuous_centered) * block_centered | -0.00 | -0.01 – 0.01 | 0.936 |
| (TrialType2 * BlockType_continuous_centered) * block_centered | -0.00 | -0.01 – 0.01 | 0.940 |
| (target [PFC_cTBS] * TrialType2 * BlockType_continuous_centered) * block_centered | 0.00 | -0.01 – 0.02 | 0.614 |
| (target [PFC_iTBS] * TrialType2 * BlockType_continuous_centered) * block_centered | 0.01 | -0.01 – 0.02 | 0.263 |
| **Random Effects** |  |  |  |
| $\sigma^2$ | | | 0.01 |
| $\tau_{00}$ subject | | | 0.00 |
| ICC |  |  | 0.01 |
| $N_{\text{subject}}$ | | | 24 |
| Observations |  |  | 22391 |
| Marginal $R^2$ / Conditional $R^2$ | | | 0.004 / 0.011 |

**Table 1b.** Movement trajectory precision.

| *Predictors* | *Estimates* | *CI* | *p* |
| --- | --- | --- | --- |
| (Intercept) | 11.32 | 8.27 – 14.36 | <**0.001** |
| target [PFC_cTBS] | -0.97 | -2.50 – 0.56 | 0.212 |
| target [PFC_iTBS] | -0.75 | -2.28 – 0.79 | 0.340 |
| TrialType2 | 10.30 | 8.75 – 11.86 | <**0.001** |
| BlockType_continuous_centered | -2.66 | -7.13 – 1.82 | 0.244 |
| block_centered | 0.10 | -0.31 – 0.52 | 0.633 |
| target [PFC_cTBS] : TrialType2 | 1.96 | -0.23 – 4.14 | 0.079 |
| target [PFC_iTBS] : TrialType2 | 1.29 | -0.90 – 3.47 | 0.249 |
| target [PFC_cTBS] * BlockType_continuous_centered | 1.67 | -4.66 – 8.01 | 0.605 |
| target [PFC_iTBS] * BlockType_continuous_centered | 0.57 | -5.79 – 6.93 | 0.861 |
| TrialType2 * BlockType_continuous_centered | 10.82 | 4.44 – 17.20 | **0.001** |
| target [PFC_cTBS] * block_centered | 0.20 | -0.39 – 0.78 | 0.512 |
| target [PFC_iTBS] * block_centered | 0.11 | -0.47 – 0.69 | 0.721 |
| TrialType2 * block_centered | 0.46 | -0.13 – 1.05 | 0.126 |
| BlockType_continuous_centered * block_centered | 0.43 | -1.45 – 2.30 | 0.655 |
| (target [PFC_cTBS] * TrialType2) * BlockType_continuous_centered | -2.37 | -11.40 – 6.67 | 0.608 |
| (target [PFC_iTBS] * TrialType2) * BlockType_continuous_centered | 2.19 | -6.91 – 11.28 | 0.637 |
| (target [PFC_cTBS] * TrialType2) * block_centered | 0.14 | -0.69 – 0.97 | 0.737 |
| (target [PFC_iTBS] * TrialType2) * block_centered | 0.01 | -0.82 – 0.84 | 0.987 |
| (target [PFC_cTBS] * BlockType_continuous_centered) * block_centered | -1.69 | -4.33 – 0.95 | 0.209 |
| (target [PFC_iTBS] * BlockType_continuous_centered) * block_centered | -0.95 | -3.62 – 1.73 | 0.487 |
| (TrialType2 * BlockType_continuous_centered) * block_centered | -0.92 | -3.56 – 1.73 | 0.496 |
| (target [PFC_cTBS] * TrialType2 * BlockType_continuous_centered) * block_centered | 1.04 | -2.69 – 4.76 | 0.585 |
| (target [PFC_iTBS] * TrialType2 * BlockType_continuous_centered) * block_centered | 1.96 | -1.84 – 5.76 | 0.313 |
| **Random Effects** |  |  |  |
| $\sigma^2$ | | | 885.35 |
| $\tau_{00}$ subject | | | 50.55 |
| ICC |  |  | 0.05 |
| $N$ subject | | | 24 |
| Observations |  |  | 22125 |
| Marginal $R^2$ / Conditional $R^2$ | | | 0.033 / 0.085 |

**Table 1c.** Movement initiation.

| *Predictors* | *Estimates* | *CI* | *p* |
| --- | --- | --- | --- |
| (Intercept) | 492.91 | 472.47 – 513.36 | <**0.001** |
| target [PFC_cTBS] | -5.96 | -13.43 – 1.51 | 0.118 |
| target [PFC iTBS] | 7.87 | 0.40 – 15.35 | **0.039** |
| TrialType2 | 54.39 | 46.81 – 61.97 | <**0.001** |
| BlockType_continuous_centered | -97.73 | -119.58 – -75.89 | <**0.001** |
| block_centered | -3.25 | -5.28 – -1.21 | **0.002** |
| target [PFC_cTBS] : TrialType2 | 1.54 | -9.13 – 12.21 | 0.778 |
| target [PFC iTBS] : TrialType2 | -1.87 | -12.55 – 8.81 | 0.731 |
| target [PFC_cTBS] * BlockType_continuous_centered | 27.83 | -3.09 – 58.75 | 0.078 |
| target [PFC iTBS] * BlockType_continuous_centered | 48.11 | 17.06 – 79.15 | **0.002** |
| TrialType2 * BlockType_continuous_centered | 43.83 | 12.68 – 74.97 | **0.006** |
| target [PFC_cTBS] * block_centered | -1.82 | -4.68 – 1.04 | 0.213 |
| target [PFC iTBS] * block_centered | 2.01 | -0.82 – 4.83 | 0.165 |
| TrialType2 * block_centered | -0.58 | -3.46 – 2.30 | 0.694 |
| BlockType_continuous_centered * block_centered | 15.75 | 6.60 – 24.91 | **0.001** |
| (target [PFC_cTBS] * TrialType2) * BlockType_continuous_centered | -17.50 | -61.60 – 26.60 | 0.437 |
| (target [PFC iTBS] * TrialType2) * BlockType_continuous_centered | -26.59 | -70.98 – 17.79 | 0.240 |
| (target [PFC_cTBS] * TrialType2) * block_centered | 0.79 | -3.27 – 4.85 | 0.702 |
| (target [PFC iTBS] * TrialType2) * block_centered | -2.13 | -6.18 – 1.92 | 0.303 |
| (target [PFC_cTBS] * BlockType_continuous_centered) * block_centered | -14.82 | -27.71 – -1.94 | **0.024** |
| (target [PFC iTBS] * BlockType_continuous_centered) * block_centered | -4.24 | -17.29 – 8.81 | 0.524 |
| (TrialType2 * BlockType_continuous_centered) * block_centered | -1.80 | -14.71 – 11.10 | 0.784 |
| (target [PFC_cTBS] * TrialType2 * BlockType_continuous_centered) * block_centered | 16.98 | -1.20 – 35.17 | 0.067 |
| (target [PFC iTBS] * TrialType2 * BlockType_continuous_centered) * block_centered | -9.95 | -28.50 – 8.60 | 0.293 |
| **Random Effects** |  |  |  |
| $\sigma^2$ | | | 21098.49 |
| $\tau_{00}$ subject | | | 2435.66 |
| ICC |  |  | 0.10 |
| N<sub>subject</sub> |  |  | 24 |
| Observations |  |  | 22125 |
| Marginal R<sup>2</sup> / Conditional R<sup>2</sup> |  |  | 0.058 / 0.156 |

**Table 1d.**
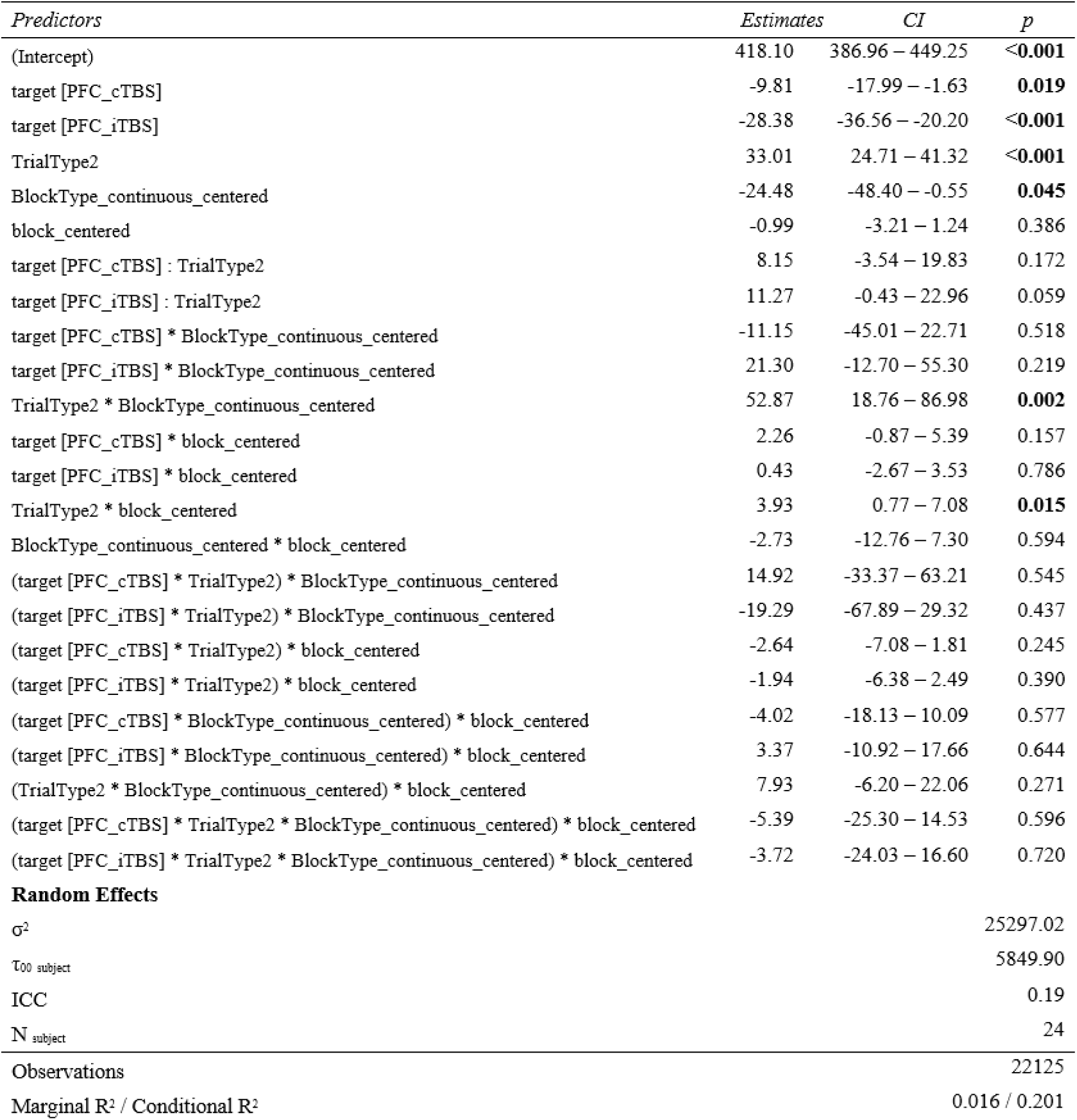
Movement duration.

**Table 1e.**
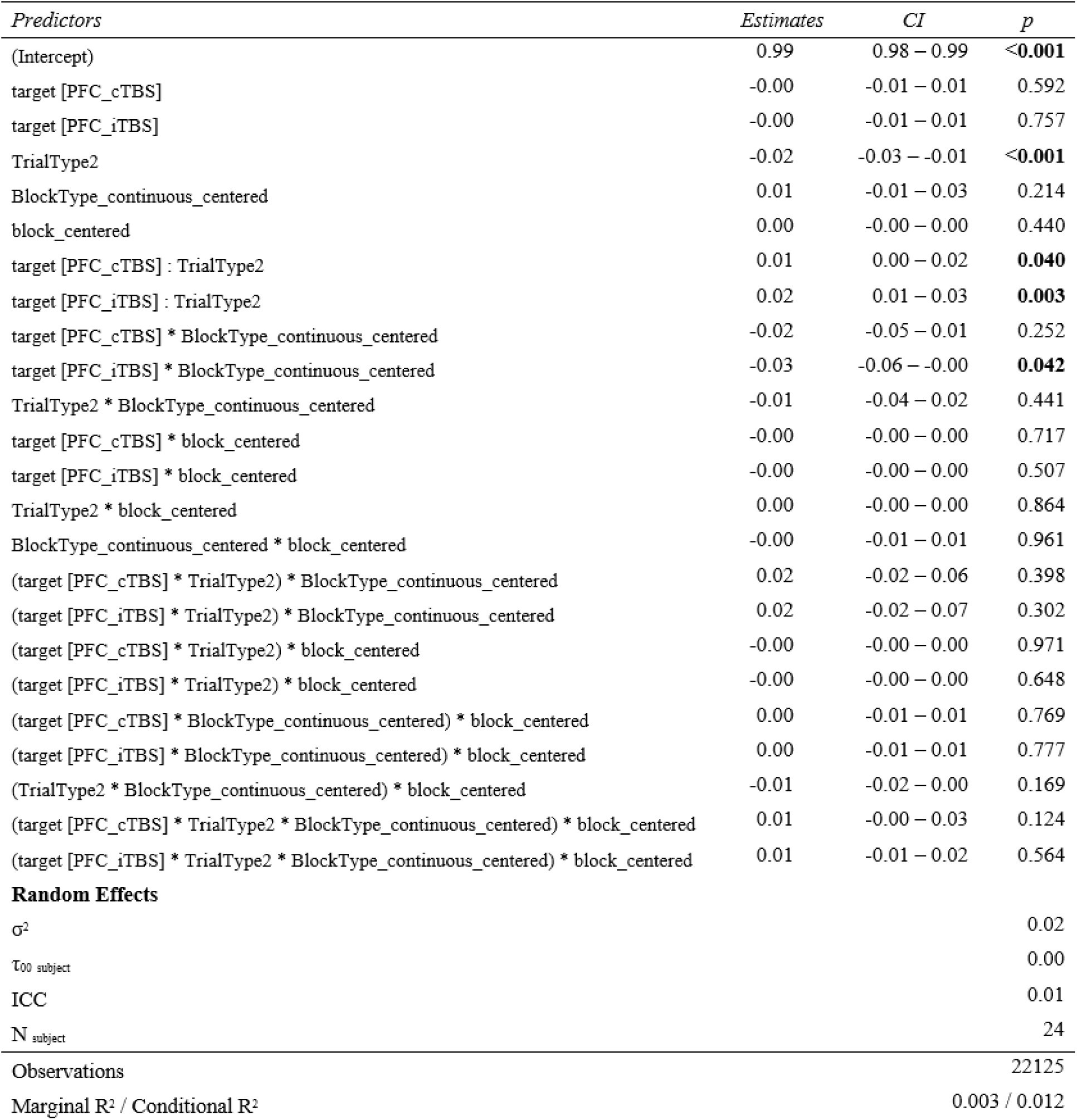
WM probe accuracy.

| *Predictors* | *Estimates* | *CI* | *p* |
| --- | --- | --- | --- |
| (Intercept) | 0.99 | 0.98 – 0.99 | <**0.001** |
| target [PFC_cTBS] | -0.00 | -0.01 – 0.01 | 0.592 |
| target [PFC_iTBS] | -0.00 | -0.01 – 0.01 | 0.757 |
| TrialType2 | -0.02 | -0.03 – -0.01 | <**0.001** |
| BlockType_continuous_centered | 0.01 | -0.01 – 0.03 | 0.214 |
| block_centered | 0.00 | -0.00 – 0.00 | 0.440 |
| target [PFC_cTBS] : TrialType2 | 0.01 | 0.00 – 0.02 | **0.040** |
| target [PFC_iTBS] : TrialType2 | 0.02 | 0.01 – 0.03 | **0.003** |
| target [PFC_cTBS] * BlockType_continuous_centered | -0.02 | -0.05 – 0.01 | 0.252 |
| target [PFC_iTBS] * BlockType_continuous_centered | -0.03 | -0.06 – -0.00 | **0.042** |
| TrialType2 * BlockType_continuous_centered | -0.01 | -0.04 – 0.02 | 0.441 |
| target [PFC_cTBS] * block_centered | -0.00 | -0.00 – 0.00 | 0.717 |
| target [PFC_iTBS] * block_centered | -0.00 | -0.00 – 0.00 | 0.507 |
| TrialType2 * block_centered | 0.00 | -0.00 – 0.00 | 0.864 |
| BlockType_continuous_centered * block_centered | -0.00 | -0.01 – 0.01 | 0.961 |
| (target [PFC_cTBS] * TrialType2) * BlockType_continuous_centered | 0.02 | -0.02 – 0.06 | 0.398 |
| (target [PFC_iTBS] * TrialType2) * BlockType_continuous_centered | 0.02 | -0.02 – 0.07 | 0.302 |
| (target [PFC_cTBS] * TrialType2) * block_centered | -0.00 | -0.00 – 0.00 | 0.971 |
| (target [PFC_iTBS] * TrialType2) * block_centered | -0.00 | -0.00 – 0.00 | 0.648 |
| (target [PFC_cTBS] * BlockType_continuous_centered) * block_centered | 0.00 | -0.01 – 0.01 | 0.769 |
| (target [PFC_iTBS] * BlockType_continuous_centered) * block_centered | 0.00 | -0.01 – 0.01 | 0.777 |
| (TrialType2 * BlockType_continuous_centered) * block_centered | -0.01 | -0.02 – 0.00 | 0.169 |
| (target [PFC_cTBS] * TrialType2 * BlockType_continuous_centered) * block_centered | 0.01 | -0.00 – 0.03 | 0.124 |
| (target [PFC_iTBS] * TrialType2 * BlockType_continuous_centered) * block_centered | 0.01 | -0.01 – 0.02 | 0.564 |
| **Random Effects** |  |  |  |
| $\sigma^2$ | | | 0.02 |
| $\tau_{00}$ subject | | | 0.00 |
| ICC |  |  | 0.01 |
| N<sub>subject</sub> |  |  | 24 |
| Observations |  |  | 22125 |
| Marginal R<sup>2</sup> / Conditional R<sup>2</sup> |  |  | 0.003 / 0.012 |

**Table 1f.** WM probe RT.

| *Predictors* | *Estimates* | *CI* | *p* |
| --- | --- | --- | --- |
| (Intercept) | 697.65 | 658.41 – 736.89 | <0.001 |
| target [PFC_cTBS] | -5.43 | -17.05 – 6.19 | 0.359 |
| target [PFC_iTBS] | -13.47 | -25.10 – -1.85 | **0.023** |
| TrialType2 | 18.28 | 6.49 – 30.08 | **0.002** |
| BlockType_continuous_centered | -9.11 | -43.11 – 24.88 | 0.599 |
| block_centered | -4.00 | -7.16 – -0.84 | **0.013** |
| target [PFC_cTBS] : TrialType2 | -1.95 | -18.55 – 14.65 | 0.818 |
| target [PFC_iTBS] : TrialType2 | 2.09 | -14.53 – 18.70 | 0.806 |
| target [PFC_cTBS] * BlockType_continuous_centered | 13.39 | -34.72 – 61.50 | 0.586 |
| target [PFC_iTBS] * BlockType_continuous_centered | 18.50 | -29.80 – 66.81 | 0.453 |
| TrialType2 * BlockType_continuous_centered | 18.48 | -29.98 – 66.94 | 0.455 |
| target [PFC_cTBS] * block_centered | 3.02 | -1.43 – 7.47 | 0.184 |
| target [PFC_iTBS] * block_centered | 1.73 | -2.67 – 6.13 | 0.441 |
| TrialType2 * block_centered | 2.92 | -1.56 – 7.40 | 0.202 |
| BlockType_continuous_centered * block_centered | 15.55 | 1.30 – 29.80 | **0.032** |
| (target [PFC_cTBS] * TrialType2) * BlockType_continuous_centered | -24.06 | -92.67 – 44.56 | 0.492 |
| (target [PFC_iTBS] * TrialType2) * BlockType_continuous_centered | -18.83 | -87.89 – 50.23 | 0.593 |
| (target [PFC_cTBS] * TrialType2) * block_centered | -4.94 | -11.25 – 1.38 | 0.125 |
| (target [PFC_iTBS] * TrialType2) * block_centered | -4.33 | -10.64 – 1.97 | 0.178 |
| (target [PFC_cTBS] * BlockType_continuous_centered) * block_centered | -19.14 | -39.19 – 0.91 | 0.061 |
| (target [PFC_iTBS] * BlockType_continuous_centered) * block_centered | -8.02 | -28.32 – 12.28 | 0.439 |
| (TrialType2 * BlockType_continuous_centered) * block_centered | 2.48 | -17.61 – 22.56 | 0.809 |
| (target [PFC_cTBS] * TrialType2 * BlockType_continuous_centered) * block_centered | 5.22 | -23.07 – 33.51 | 0.718 |
| (target [PFC_iTBS] * TrialType2 * BlockType_continuous_centered) * block_centered | 3.94 | -24.93 – 32.80 | 0.789 |
| **Random Effects** |  |  |  |
| $\sigma^2$ | | | 51071.79 |
| $\tau_{00}$ subject | | | 9195.98 |
| ICC |  |  | 0.15 |
| N<sub>subject</sub> |  |  | 24 |
| Observations |  |  | 22125 |
| Marginal R<sup>2</sup> / Conditional R<sup>2</sup> |  |  | 0.003 / 0.155 |

### Two-target models (cTBS vs. iTBS)

**Table 2a.** Movement choice accuracy.

**Two-target models (cTBS vs. iTBS)****Table 2a. Movement choice accuracy**
| *Predictors* | *Estimates* | *CI* | *p* |
| --- | --- | --- | --- |
| (Intercept) | 0.99 | 0.99 – 1.00 | <**0.001** |
| target [PFC_iTBS] | 0.00 | -0.00 – 0.01 | 0.932 |
| TrialType [incompatible] | -0.01 | -0.02 – -0.01 | <**0.001** |
| BlockType_continuous_centered | 0.00 | -0.01 – 0.02 | 0.619 |
| block_centered | -0.00 | -0.00 – -0.00 | **0.041** |
| target [PFC_iTBS] * TrialType [incompatible] | 0.00 | -0.01 – 0.01 | 0.732 |
| target [PFC_iTBS] * BlockType_continuous_centered | 0.01 | -0.01 – 0.03 | 0.398 |
| TrialType [incompatible] * BlockType_continuous_centered | -0.03 | -0.05 – -0.01 | **0.013** |
| target [PFC_iTBS] * block_centered | 0.00 | -0.00 – 0.00 | 0.087 |
| TrialType [incompatible] * block_centered | 0.00 | 0.00 – 0.00 | **0.006** |
| BlockType_continuous_centered * block_centered | 0.00 | -0.00 – 0.01 | 0.442 |
| (target [PFC_iTBS] * TrialType [incompatible]) * BlockType_continuous_centered | 0.00 | -0.03 – 0.03 | 0.789 |
| (target [PFC_iTBS] * TrialType [incompatible]) * block_centered | -0.00 | -0.01 – -0.00 | **0.049** |
| (target [PFC_iTBS] * BlockType_continuous_centered) * block_centered | -0.00 | -0.01 – 0.01 | 0.452 |
| (TrialType [incompatible] * BlockType_continuous_centered) * block_centered | 0.00 | -0.01 – 0.01 | 0.509 |
| (target [PFC_iTBS] * TrialType [incompatible] * BlockType_continuous_centered) * block_centered | 0.00 | -0.01 – 0.02 | 0.545 |
| **Random Effects** |  |  |  |
| $\sigma^2$ | | | 0.01 |
| $\tau_{00 \text{ subject}}$ | | | 0.00 |
| ICC |  |  | 0.01 |
| $N_{\text{subject}}$ | | | 24 |
| Observations |  |  | 14974 |
| Marginal $R^2$ / Conditional $R^2$ | | | 0.004 / 0.012 |

**Table 2b.**
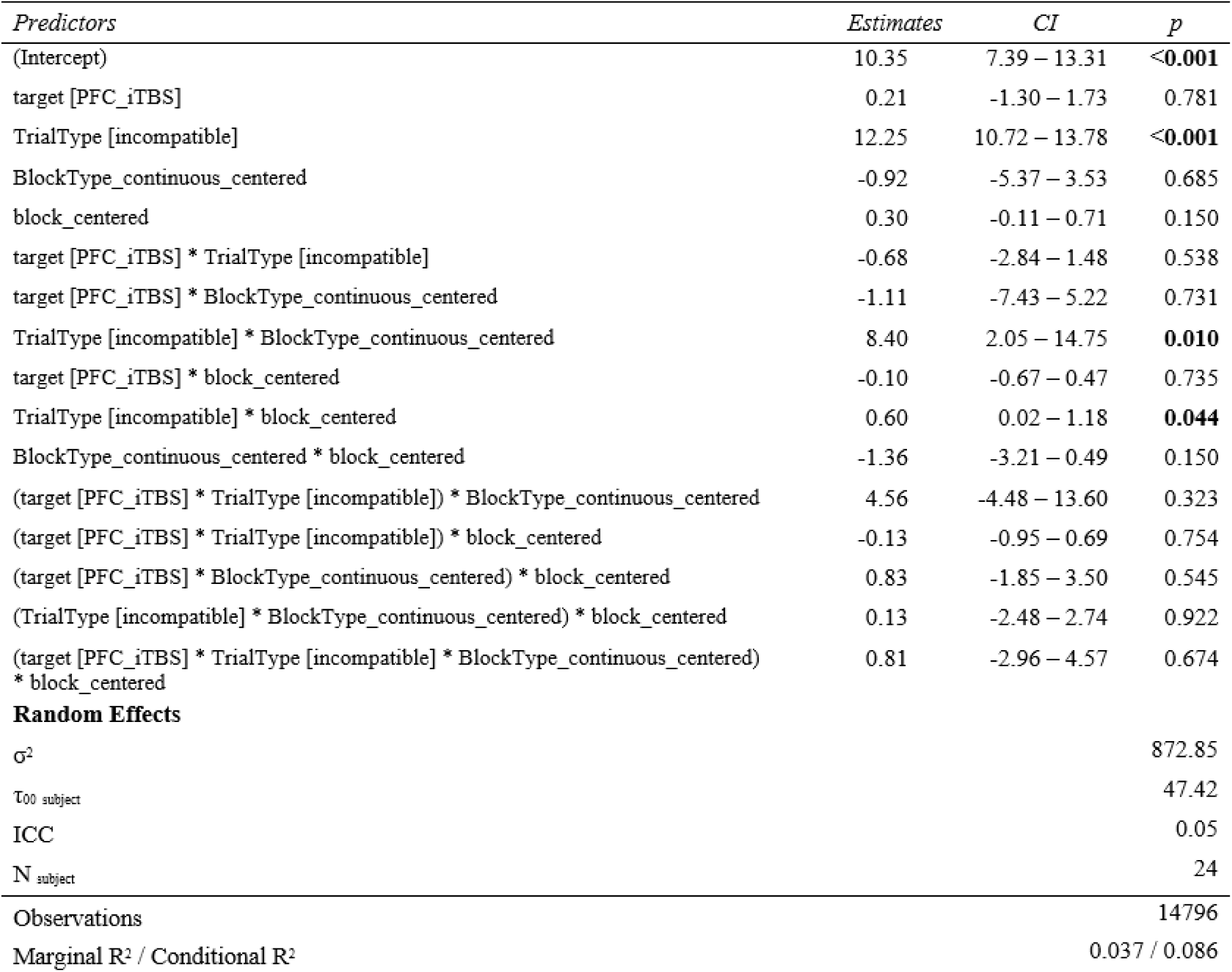
Movement trajectory precision.

**Table 2c.** Movement initiation.

| *Predictors* | *Estimates* | *CI* | *p* |
| --- | --- | --- | --- |
| (Intercept) | 487.07 | 464.51 – 509.63 | <0.001 |
| target [PFC iTBS] | 13.81 | 6.40 – 21.22 | <0.001 |
| TrialType [incompatible] | 56.00 | 48.52 – 63.48 | <0.001 |
| BlockType_continuous_centered | -68.82 | -90.62 – -47.03 | <0.001 |
| block_centered | -5.06 | -7.06 – -3.05 | <0.001 |
| target [PFC iTBS] * TrialType [incompatible] | -3.55 | -14.14 – 7.03 | 0.511 |
| target [PFC iTBS] * BlockType_continuous_centered | 18.75 | -12.21 – 49.71 | 0.235 |
| TrialType [incompatible] * BlockType_continuous_centered | 26.49 | -4.60 – 57.59 | 0.095 |
| target [PFC iTBS] * block_centered | 3.83 | 1.03 – 6.63 | 0.007 |
| TrialType [incompatible] * block_centered | 0.28 | -2.57 – 3.13 | 0.846 |
| BlockType_continuous_centered * block_centered | -1.17 | -10.23 – 7.89 | 0.801 |
| (target [PFC iTBS] * TrialType [incompatible]) * BlockType_continuous_centered | -9.85 | -54.11 – 34.40 | 0.663 |
| (target [PFC iTBS] * TrialType [incompatible]) * block_centered | -2.94 | -6.97 – 1.08 | 0.152 |
| (target [PFC iTBS] * BlockType_continuous_centered) * block_centered | 13.61 | 0.50 – 26.71 | 0.042 |
| (TrialType [incompatible] * BlockType_continuous_centered) * block_centered | 14.77 | 2.00 – 27.54 | 0.023 |
| (target [PFC iTBS] * TrialType [incompatible] * BlockType_continuous_centered) * block_centered | -26.39 | -44.83 – -7.95 | 0.005 |
| **Random Effects** |  |  |  |
| $\sigma^2$ | | | 20920.12 |
| $\tau_{00}$ subject | | | 3007.24 |
| ICC |  |  | 0.13 |
| $N$ subject | | | 24 |
| Observations |  |  | 14796 |
| Marginal $R^2$ / Conditional $R^2$ | | | 0.053 / 0.172 |

**Table 2d.** Movement duration.

| *Predictors* | *Estimates* | *CI* | *p* |
| --- | --- | --- | --- |
| (Intercept) | 408.25 | 375.17 – 441.32 | <0.001 |
| target [PFC_iTBS] | -18.58 | -26.70 – -10.47 | <0.001 |
| TrialType [incompatible] | 41.31 | 33.12 – 49.50 | <0.001 |
| BlockType_continuous_centered | -34.48 | -58.35 – -10.61 | 0.005 |
| block_centered | 1.32 | -0.88 – 3.51 | 0.239 |
| target [PFC_iTBS] * TrialType [incompatible] | 2.62 | -8.97 – 14.21 | 0.658 |
| target [PFC_iTBS] * BlockType_continuous_centered | 32.09 | -1.83 – 66.00 | 0.064 |
| TrialType [incompatible] * BlockType_continuous_centered | 67.12 | 33.06 – 101.17 | <0.001 |
| target [PFC_iTBS] * block_centered | -1.97 | -5.04 – 1.10 | 0.209 |
| TrialType [incompatible] * block_centered | 1.28 | -1.84 – 4.41 | 0.420 |
| BlockType_continuous_centered * block_centered | -8.13 | -18.05 – 1.80 | 0.108 |
| (target [PFC_iTBS] * TrialType [incompatible]) * BlockType_continuous_centered | -33.12 | -81.59 – 15.36 | 0.181 |
| (target [PFC_iTBS] * TrialType [incompatible]) * block_centered | 0.63 | -3.78 – 5.03 | 0.781 |
| (target [PFC_iTBS] * BlockType_continuous_centered) * block_centered | 8.33 | -6.03 – 22.69 | 0.255 |
| (TrialType [incompatible] * BlockType_continuous_centered) * block_centered | 2.83 | -11.16 – 16.81 | 0.692 |
| (target [PFC_iTBS] * TrialType [incompatible] * BlockType_continuous_centered) * block_centered | -1.28 | -21.48 – 18.92 | 0.901 |
| **Random Effects** |  |  |  |
| $\sigma^2$ | | | 25097.70 |
| $\tau_{00}$ subject | | | 6630.64 |
| ICC |  |  | 0.21 |
| N<sub>subject</sub> |  |  | 24 |
| Observations |  |  | 14796 |
| Marginal R<sup>2</sup> / Conditional R<sup>2</sup> |  |  | 0.017 / 0.222 |

**Table 2e.** WM probe accuracy.

| *Predictors* | *Estimates* | *CI* | *p* |
| --- | --- | --- | --- |
| (Intercept) | 0.98 | 0.98 – 0.99 | <**0.001** |
| target [PFC iTBS] | 0.00 | -0.01 – 0.01 | 0.815 |
| TrialType [incompatible] | -0.01 | -0.01 – -0.00 | **0.045** |
| BlockType_continuous_centered | -0.00 | -0.02 – 0.02 | 0.713 |
| block_centered | 0.00 | -0.00 – 0.00 | 0.766 |
| target [PFC iTBS] * TrialType [incompatible] | 0.00 | -0.01 – 0.01 | 0.336 |
| target [PFC iTBS] * BlockType_continuous_centered | -0.01 | -0.04 – 0.02 | 0.346 |
| TrialType [incompatible] * BlockType_continuous_centered | 0.01 | -0.02 – 0.04 | 0.671 |
| target [PFC iTBS] * block_centered | -0.00 | -0.00 – 0.00 | 0.744 |
| TrialType [incompatible] * block_centered | 0.00 | -0.00 – 0.00 | 0.903 |
| BlockType_continuous_centered * block_centered | 0.00 | -0.01 – 0.01 | 0.800 |
| (target [PFC iTBS] * TrialType [incompatible]) * BlockType_continuous_centered | 0.00 | -0.04 – 0.05 | 0.833 |
| (target [PFC iTBS] * TrialType [incompatible]) * block_centered | -0.00 | -0.00 – 0.00 | 0.663 |
| (target [PFC iTBS] * BlockType_continuous_centered) * block_centered | 0.00 | -0.01 – 0.01 | 0.934 |
| (TrialType [incompatible] * BlockType_continuous_centered) * block_centered | 0.01 | -0.01 – 0.02 | 0.373 |
| (target [PFC iTBS] * TrialType [incompatible] * BlockType_continuous_centered) * block_centered | -0.01 | -0.03 – 0.01 | 0.309 |
| **Random Effects** |  |  |  |
| $\sigma^2$ | | | 0.02 |
| $\tau_{00}$ subject | | | 0.00 |
| ICC |  |  | 0.01 |
| N<sub>subject</sub> |  |  | 24 |
| Observations |  |  | 14796 |
| Marginal R<sup>2</sup> / Conditional R<sup>2</sup> |  |  | 0.001 / 0.009 |

**Table 2f.** WM probe RT.

| *Predictors* | *Estimates* | *CI* | *p* |
| --- | --- | --- | --- |
| (Intercept) | 692.31 | 648.68 – 735.95 | <0.001 |
| target [PFC iTBS] | -8.10 | -19.67 – 3.47 | 0.170 |
| TrialType [incompatible] | 16.59 | 4.92 – 28.27 | 0.005 |
| BlockType_continuous_centered | 4.66 | -29.38 – 38.70 | 0.788 |
| block_centered | -1.02 | -4.15 – 2.11 | 0.521 |
| target [PFC iTBS] * TrialType [incompatible] | 3.49 | -13.04 – 20.02 | 0.679 |
| target [PFC iTBS] * BlockType_continuous_centered | 5.53 | -42.82 – 53.89 | 0.823 |
| TrialType [incompatible] * BlockType_continuous_centered | -5.90 | -54.46 – 42.66 | 0.812 |
| target [PFC iTBS] * block_centered | -1.31 | -5.68 – 3.07 | 0.559 |
| TrialType [incompatible] * block_centered | -1.85 | -6.30 – 2.61 | 0.417 |
| BlockType_continuous_centered * block_centered | -3.03 | -17.18 – 11.12 | 0.675 |
| (target [PFC iTBS] * TrialType [incompatible]) * BlockType_continuous_centered | 5.00 | -64.12 – 74.12 | 0.887 |
| (target [PFC iTBS] * TrialType [incompatible]) * block_centered | 0.58 | -5.70 – 6.86 | 0.856 |
| (target [PFC iTBS] * BlockType_continuous_centered) * block_centered | 9.76 | -10.71 – 30.23 | 0.350 |
| (TrialType [incompatible] * BlockType_continuous_centered) * block_centered | 7.58 | -12.36 – 27.52 | 0.456 |
| (target [PFC iTBS] * TrialType [incompatible] * BlockType_continuous_centered) * block_centered | -1.19 | -29.99 – 27.61 | 0.935 |
| **Random Effects** |  |  |  |
| $\sigma^2$ | | | 51028.95 |
| $\tau_{00}$ subject | | | 11478.95 |
| ICC |  |  | 0.18 |
| $N$ subject | | | 24 |
| Observations |  |  | 14796 |
| Marginal $R^2$ / Conditional $R^2$ | | | 0.003 / 0.186 |

### Trial history models (current x previous trial compatibility)

**Table 3a.**
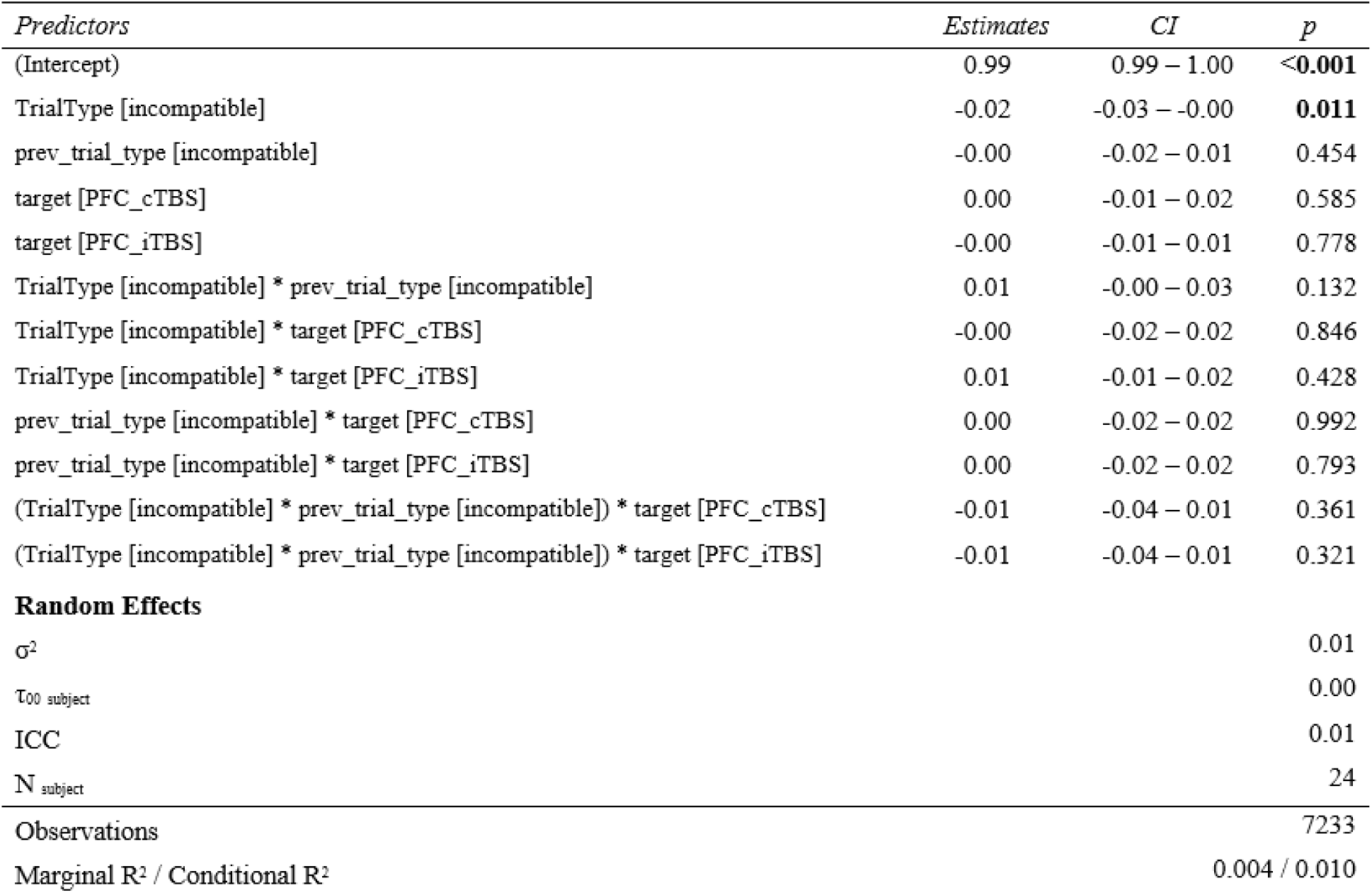
Movement choice accuracy.

**Table 3b.** Movement trajectory precision.

| *Predictors* | *Estimates* | *CI* | *p* |
| --- | --- | --- | --- |
| (Intercept) | 10.66 | 6.68 – 14.64 | <0.001 |
| TrialType [incompatible] | 9.76 | 6.35 – 13.16 | <0.001 |
| prev_trial_type [incompatible] | 1.61 | -1.77 – 5.00 | 0.350 |
| target [PFC_cTBS] | 0.04 | -3.35 – 3.44 | 0.980 |
| target [PFC_iTBS] | -1.31 | -4.75 – 2.12 | 0.454 |
| TrialType [incompatible] * prev_trial_type [incompatible] | -2.49 | -7.32 – 2.35 | 0.314 |
| TrialType [incompatible] * target [PFC_cTBS] | 1.47 | -3.32 – 6.26 | 0.548 |
| TrialType [incompatible] * target [PFC_iTBS] | 2.40 | -2.41 – 7.21 | 0.329 |
| prev_trial_type [incompatible] * target [PFC_cTBS] | -1.56 | -6.31 – 3.19 | 0.521 |
| prev_trial_type [incompatible] * target [PFC_iTBS] | -0.15 | -4.92 – 4.62 | 0.950 |
| (TrialType [incompatible] * prev_trial_type [incompatible]) * target [PFC_cTBS] | 1.77 | -5.04 – 8.57 | 0.611 |
| (TrialType [incompatible] * prev_trial_type [incompatible]) * target [PFC_iTBS] | -1.87 | -8.69 – 4.94 | 0.590 |
| **Random Effects** |  |  |  |
| $\sigma^2$ | | | 889.65 |
| $\tau_{00}$ subject | | | 62.03 |
| ICC |  |  | 0.07 |
| $N$ subject | | | 24 |
| Observations |  |  | 7137 |
| Marginal $R^2$ / Conditional $R^2$ | | | 0.026 / 0.089 |

**Table 3c.**
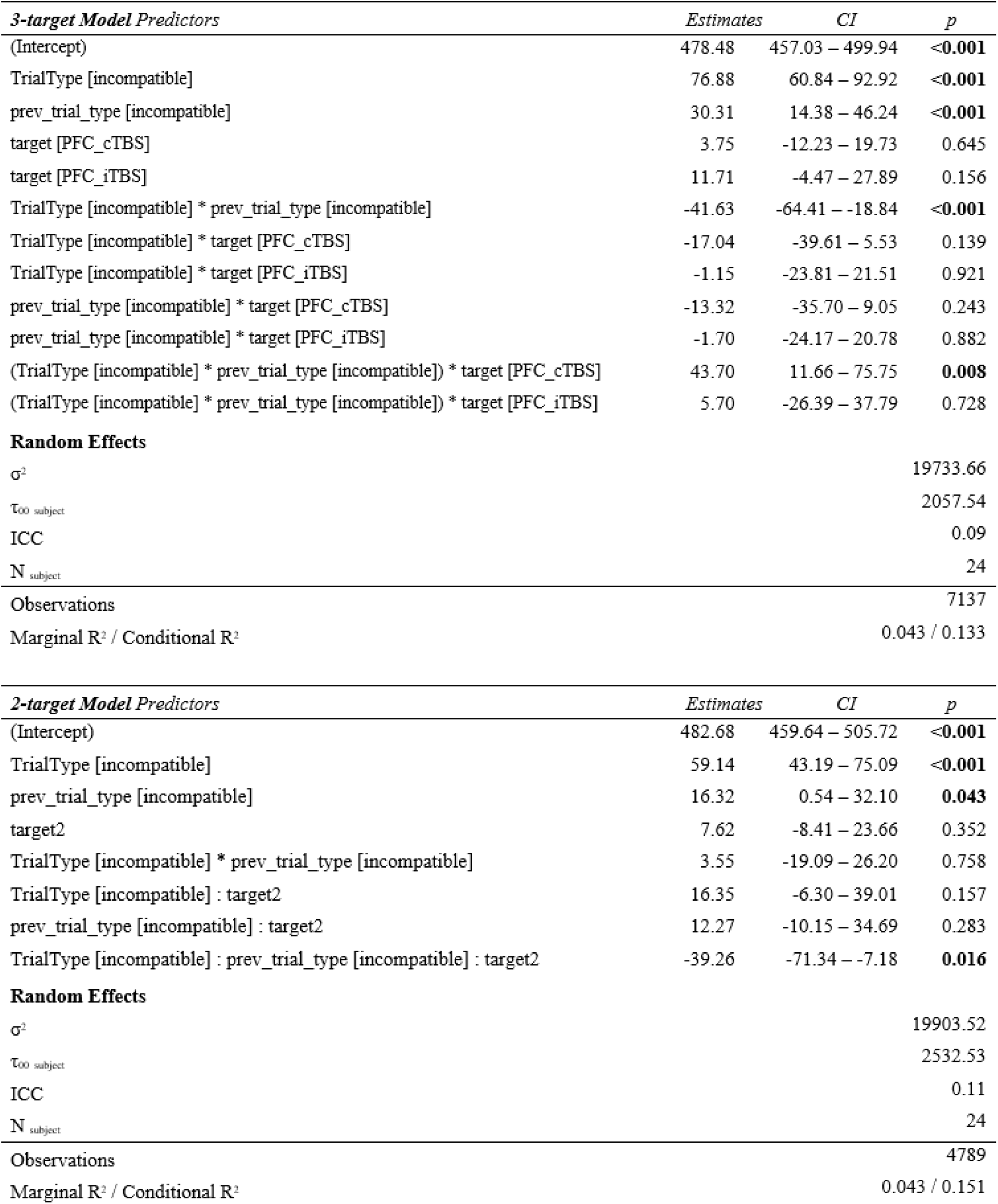
Movement initiation.

**Table 3d.** Movement duration.

| *Predictors* | *Estimates* | *CI* | *p* |
| --- | --- | --- | --- |
| (Intercept) | 404.21 | 371.34 – 437.08 | <0.001 |
| TrialType [incompatible] | 42.53 | 24.14 – 60.91 | <0.001 |
| prev_trial_type [incompatible] | 15.41 | -2.86 – 33.67 | 0.098 |
| target [PFC_cTBS] | 8.22 | -10.10 – 26.54 | 0.379 |
| target [PFC_iTBS] | -17.20 | -35.75 – 1.34 | 0.069 |
| TrialType [incompatible] * prev_trial_type [incompatible] | -25.00 | -51.12 – 1.12 | 0.061 |
| TrialType [incompatible] * target [PFC_cTBS] | 5.95 | -19.93 – 31.82 | 0.652 |
| TrialType [incompatible] * target [PFC_iTBS] | 6.81 | -19.16 – 32.78 | 0.607 |
| prev_trial_type [incompatible] * target [PFC_cTBS] | -6.79 | -32.44 – 18.86 | 0.604 |
| prev_trial_type [incompatible] * target [PFC_iTBS] | -6.83 | -32.59 – 18.93 | 0.603 |
| (TrialType [incompatible] * prev_trial_type [incompatible]) * target [PFC_cTBS] | -5.74 | -42.48 – 30.99 | 0.759 |
| (TrialType [incompatible] * prev_trial_type [incompatible]) * target [PFC_iTBS] | 10.08 | -26.70 – 46.87 | 0.591 |
| **Random Effects** |  |  |  |
| $\sigma^2$ | | | 25930.95 |
| $\tau_{00}$ subject | | | 5673.64 |
| ICC |  |  | 0.18 |
| N<sub>subject</sub> |  |  | 24 |
| Observations |  |  | 7137 |
| Marginal R<sup>2</sup> / Conditional R<sup>2</sup> |  |  | 0.013 / 0.191 |

